# A genome-scale screen identifies sulfated glycosaminoglycans as pivotal in epithelial cell damage by *Candida albicans*

**DOI:** 10.1101/2024.05.23.595417

**Authors:** Jianfeng Lin, Jian Miao, Katherine G. Schaefer, Charles M. Russell, Robert J. Pyron, Fuming Zhang, Quynh T. Phan, Norma V. Solis-Swidergall, Hong Liu, Masato Tashiro, Jonathan S. Dordick, Robert J. Linhardt, Michael R. Yeaman, Gavin M. King, Francisco N. Barrera, Brian M. Peters, Scott G. Filler

## Abstract

Candidalysin is a cytolytic peptide produced by the opportunistic fungal pathogen *Candida albicans.* This peptide is a key virulence factor in mouse models of mucosal and hematogenously disseminated candidiasis. Despite intense interest in the role of candidalysin in *C. albicans* pathogenicity, its host cell targets have remained elusive. To fill this knowledge gap, we performed a genome-wide loss-of-function CRISPR screen in a human oral epithelial cell line to identify specific host factors required for susceptibility to candidalysin-induced cellular damage. Among the top hits were *XYLT2*, *B3GALT6* and *B3GAT3*, genes that function in glycosaminoglycan (GAG) biosynthesis. Deletion of these genes led to the absence of GAGs such as heparan sulfate on the epithelial cell surface and increased resistance to damage induced by both candidalysin and live *C. albicans.* Biophysical analyses including surface plasmon resonance and atomic force and electron microscopy indicated that candidalysin physically binds to sulfated GAGs, facilitating its oligomerization or enrichment on the host cell surface. The addition of exogenous sulfated GAGs or the GAG analogue dextran sulfate protected cells against candidalysin-induced damage. Dextran sulfate, but not non-sulfated dextran, also inhibited epithelial cell endocytosis of *C. albicans* and fungal-induced epithelial cell cytokine and chemokine production. In a murine model of vulvovaginal candidiasis, topical dextran sulfate administration reduced host tissue damage and decreased intravaginal IL-1β and neutrophil levels. Collectively, these data indicate that GAGs are epithelial cell targets of candidalysin and can be used therapeutically to protect cells from candidalysin-induced damage.

## Introduction

*Candida albicans*, a commensal fungus, frequently colonizes epithelial surfaces of healthy individuals including the skin and oral, gut, and vaginal mucosa. When local or systemic host defenses are dysregulated, this opportunistic fungus can become pathogenic, causing mucosal infections such as oropharyngeal and vulvovaginal candidiasis, or life-threatening invasive infections such as hematogenously disseminated candidiasis.

A key virulence factor of *C. albicans* is candidalysin, a cytolytic peptide toxin required for maximal virulence in mouse models of mucosal and hematogenously disseminated candidiasis ^1–3^. Candidalysin is synthesized by the fungus as a pro-protein encoded by the gene *ECE1.* The Kex2 protease cleaves the Ece1 pro-protein into eight peptides. One of these derivative peptides is candidalysin, a 31-amino acid, amphiphilic, a-helical oligopeptide that is released from *C. albicans* hyphae ^1,4,5^. When a *C. albicans* hypha invades a host cell, an invasion pocket is formed and candidalysin accumulates at high concentrations within this pocket ^6–8^. Candidalysin subsequently undergoes a complex polymerization process that leads to the formation of pore-like structures on the host cell membrane that cause loss of membrane integrity and an influx of calcium ions ^4,7,9,10^. This toxin stimulates an innate epithelial immune response by activating the epidermal growth factor receptor (EGFR) ^3,11^, the mitogen-activated protein kinases P38 and extracellular regulated kinases (ERK)1/2 ^12–15^, and the c-Fos transcription factor ^12,14^. These signaling pathways trigger the production of downstream inflammatory mediators such as IL-6, GM-CSF, CXCL8 and IL-1β, which recruit phagocytes to foci of infection and enhance their fungicidal activities ^2,3,11,16,17^.

Although the mechanism of action of candidalysin has been studied extensively ^2,4,5,7,8,12^, the host determinants that target the binding and oligomerization of this toxin on the host cell surface remain unknown. In this study, we performed a genome-wide loss-of-function CRISPR screen in a human oral epithelial cell line to identify specific host factors required for susceptibility to candidalysin-induced cellular damage. We found that deletion of genes involved in glycosaminoglycan (GAG) biosynthesis - namely *XYLT2*, *B3GALT6*, and *B3GAT3* - lead to the absence of GAGs such as heparan sulfate on the epithelial cell surface. The absence of these GAGs results in enhanced resistance to damage induced by both candidalysin and live *C. albicans.* Biophysical analyses indicated that candidalysin directly binds to sulfated GAGs, which facilitate its oligomerization or enrichment on the cell surface. Exogenous GAGs or GAG analogs, such as dextran sulfate, bind to candidalysin and inhibit its activity. In a mouse model of vulvovaginal candidiasis, intravaginal administration of dextran sulfate significantly reduces epithelial cell damage, IL-1β release and neutrophil accumulation. Collectively, these data indicate that host GAGs facilitate candidalysin activity and that GAG analogs can be used therapeutically to protect host cells from candidalysin-induced damage.

## Results

### A CRISPR screen identifies host factors for candidalysin-induced cell damage

To identify potential host cell targets of candidalysin, we performed a genome-wide CRISPR-Cas9 mediated knockout screen in the TR146 oral epithelial cell line. We first generated a TR146 cell line that stably expressed Cas9 (Supplementary Fig. 1a, b), and then transduced this cell line with a lentiviral sgRNA library (Brunello) that has four sgRNAs for each of 19,114 human genes ^18,19^. The transduced cells were incubated with 30 µM candidalysin for 8 h, which resulted in 60-70% cell death in control TR146 cells, as measured by a 2,3-bis-(2-methoxy-4-nitro-5-sulfophenyl)- 2h-tetrazolium-5-carboxanilide (XTT) assay (Supplementary Fig. 1c-d). After five rounds of selection with a recovery step in between each round (Fig. 1a), the genes targeted by sgRNAs in the surviving cells were identified via next-generation sequencing and the frequency of each sgRNA was compared with that in the initial library ^20^. The fold-enrichment of each targeted gene was ranked by the sigma FC score ^20^ (Fig. 1a).

**Fig. 1.**
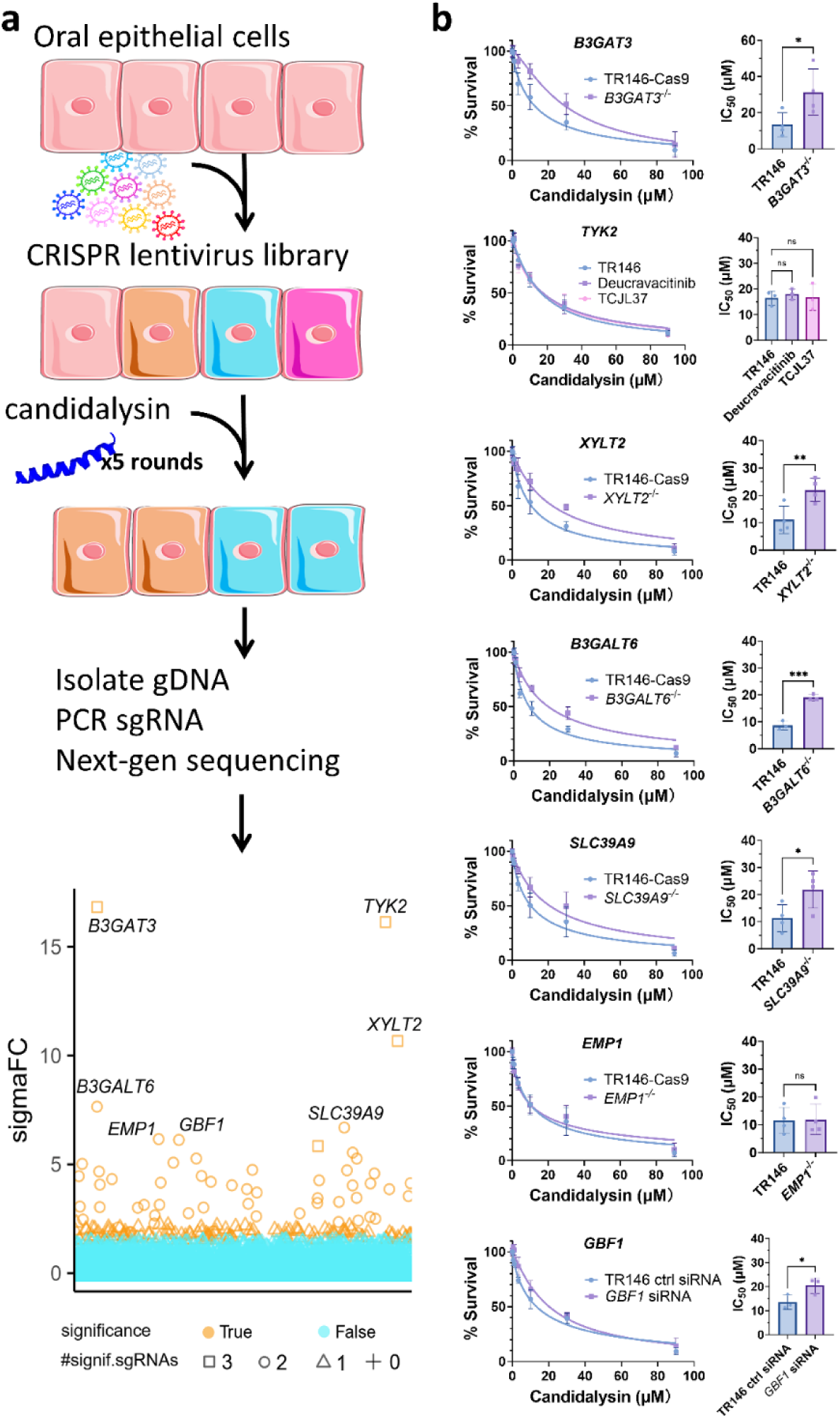
Genes identified by the genome-wide CRISPR screen that are required for maximal candidalysin-induced epithelial cell damage. **a,** Schematic diagram of the genome-wide CRISPR screen used to identify genes required for susceptibility to candidalysin-induced epithelial cell damage and scatterplot of the results. In the scatterplot, the number of sgRNAs targeting each gene that were significantly enriched is indicated with different symbols. The sigmaFC gene rank for each gene was calculated by adding the fold changes of all sgRNAs that target that gene, multiplied by the number of sgRNAs that showed significant enrichment. The top 7 enriched genes (sigmaFC > 6) are labeled in the plot. **b,** Effects of CRISPR deletion (*B3GAT3, XYLT2, B3GALT6, SLC39A9, EMP1*), pharmacologic inhibition (Tyk2), and siRNA knockdown (*GBF1*) on the survival of oral epithelial cells after 6 h of candidalysin exposure. The left panels show the survival (measured by an XTT assay) of epithelial cells exposed to the indicated concentrations of candidalysin. The plots represent the combined results of 3-4 experiments, each performed in triplicate. The right panels show the concentration of candidalysin that yielded 50% survival (IC_50_), which was calculated from the data in the corresponding graph in the left panels. Results are mean ± SD. K/O, knockout; ns, not significant; *, p<0.05; **, p<0.01; ***, p<0.001 by the unpaired, two-sided Student’s t test.

### Deletion of epithelial *XYLT2*, *B3GALT6* or *B3GAT3* confers host cell resistance to candidalysin

A total of 182 genes were significantly enriched in the screen (Supplementary Table 1). The top seven hits, based on their sigmaFC scores, were *B3GALT6, TYK2, XYLT2, B3GAT3, SLC39A9, GBF1,* and *EMP1* (Fig. 1a). To confirm that the products of these genes are indeed required for maximal damage induced by candidalysin, we used CRISPR-Cas9 gene deletion, small interfering RNA (siRNA) knockdown, or small molecule inhibitors to eliminate or reduce their activity. Each gene knockout or knockdown was confirmed via western blot analysis (Supplementary Fig. 2a). Inhibition of the Tyk2 tyrosine kinase with deucravacitinib (3 μM), TCJL37 (1 μM), or siRNA knockdown of Tyk2 had minimal effect on host cell susceptibility to candidalysin (Fig. 1b, Supplementary Fig. 2b). Deletion of *EMP1,* which encodes epithelial membrane protein 1, had no effect on host cell survival (Fig. 1b). siRNA knockdown of Gbf1, a Sec7 domain family guanine nucleotide exchange factor protein ^21^, (Supplementary Fig. 2a), resulted in slightly increased resistance to candidalysin relative to TR146 cells transfected with control siRNA. Likewise, deletion of *SLC39A9,* which is predicted to encode a zinc ion transporter ^22^, modestly, but significantly increased resistance to candidalysin-induced cell damage. Importantly, deletion of *XYLT2*, *B3GALT6* or *B3GAT3* significantly increased cell resistance to candidalysin (Fig. 1b). The IC_50_ of candidalysin in *XYLT2*^-/-^, *B3GALT6*^-/-^ or *B3GAT3*^-/-^ cells was 2.2- to 2.3-fold higher than that of the control cells. *XYLT2*, *B3GALT6* and *B3GAT3* were three of the four most significantly enriched genes, and they encode enzymes that catalyze the synthesis of the Xyl-Gal-Gal-GlcA tetrasaccharide linker that attaches GAGs to core proteins ^23–27^ (Fig. 2a). Indeed, deletion of any of the three genes markedly decreased the amount of cell-surface exposed heparan sulfate, as determined by indirect immunofluorescence and flow cytometry (Supplementary Fig. 2c, d). Based on these results, we focused our subsequent experiments on the interactions of GAGs with candidalysin.

**Fig. 2.**
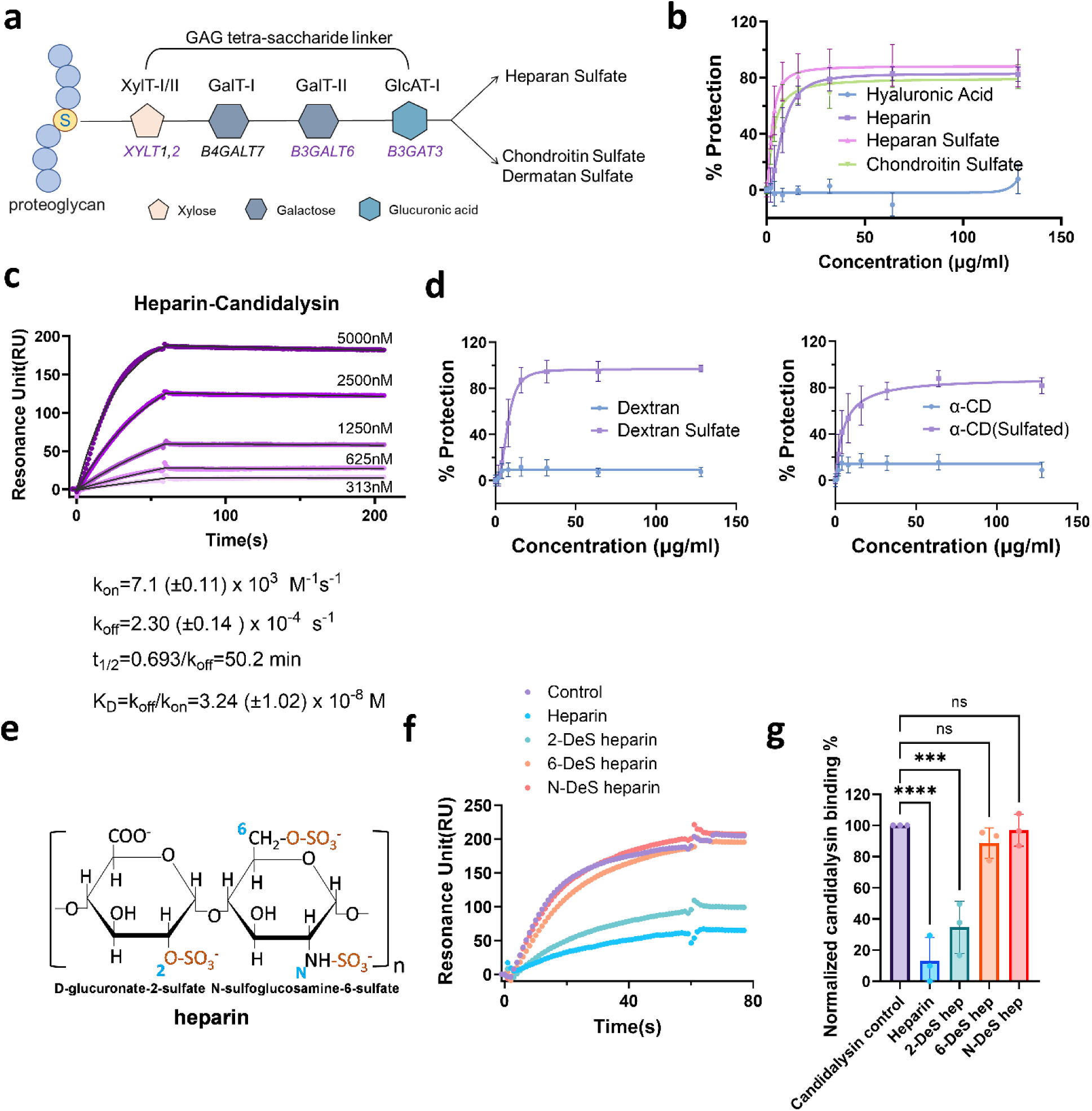
Sulfated glycosaminoglycans (GAGs) and dextrin analogs bind to candidalysin and provide dose-dependent protection to epithelial cells against candidalysin-induced damage. **a,** Diagram illustrating the enzymes that catalyze the biosynthesis of the tetra-saccharide linker present in all GAGs. The genes that encode the enzymes are denoted under the monosaccharide, and the ones in purple are those identified by the CRISPR screen. **b,** Protection from damage caused by a 6-h exposure to 30 μM candidalysin provided by the indicated GAGs as measured by an XTT assay. The curves were generated from the data of at least 3 independent experiments, each performed in triplicate. Percent protection was calculated as the normalized increase in host cell survival relative to control cells treated with candidalysin alone. **c,** Surface plasmon resonance (SPR) sensorgram of the interaction of candidalysin with heparin immobilized on the biosensor chip. The concentration of candidalysin used to generate each sensorgram is indicated on the graph. The binding kinetics (*k_on_,* association rate constant; *k_off_,* dissociation rate constant; t_1/2_, half-life; and *K_D_=k_on_/k_off_,* binding equilibrium dissociation constant) were calculated from the global fitting of the five concentrations using a 1:1 Langmuir binding model. **d,** Protection from damage caused by a 6-h exposure to 30 μM candidalysin provided by dextran, dextran sulfate, α-cyclodextrin (α-CD) or sulfated α-cyclodextrin measured by an XTT assay. The curves were generated from the data of 3 independent experiments. **e,** Structure of heparin showing the locations of the sulfate groups. **f,** Representative surface plasmon resonance sensorgrams showing the effects of 2-*O*-desulfated heparin (2-Des hep), 6-*O*-desulfated heparin (6-Des hep), and *N*-desulfated heparin (*N*-Des hep) on the interaction of candidalysin with heparin on a biosensor chip. **g,** Combined results of 3 independent experiments showing the inhibitory effects of the various desulfated heparins on the interaction of candidalysin with heparin. Results are mean ± SD. ns, not significant; ***, p<0.001; ****, p<0.0001 by one-way ANOVA with Dunnett’s multiple comparisons test.

### Candidalysin binds to GAGs

Next, we investigated whether the addition of various GAGs to the medium could compete with endogenous cell-surface GAGs and protect epithelial cells from candidalysin-induced cell damage. Oral epithelial cells were incubated with increasing concentrations of the naturally occurring GAGs, heparin, heparan sulfate, chondroitin sulfate, or hyaluronic acid (see structures in Supplementary Fig. 3a). Next, 30 μM of candidalysin was added and epithelial cell survival was measured 6 h later. Heparin, heparan sulfate, and chondroitin sulfate all exhibited dose-dependent protection against candidalysin-induced epithelial cell damage, while hyaluronic acid provided no discernible protection (Fig. 2b). These results indicate that some but not all GAGs can protect oral epithelial cells from damage caused by candidalysin, possibly by competitive binding to the toxin in the medium before it can bind to the host cells.

To test this hypothesis, we analyzed the kinetics and affinity of the candidalysin-heparin interaction using surface plasmon resonance (SPR). Different concentrations of candidalysin were passed through a sensor chip on which biotinylated heparin was immobilized. The kinetics were determined by globally fitting the association and dissociation phases, employing a 1:1 Langmuir binding model. The sensorgrams illustrated the interaction between heparin and various concentrations of candidalysin (Fig. 2c). The SPR results indicate that candidalysin binds to heparin with high affinity at nanomolar scale at pH 7.4, characterized by a K_D_ of 32.4 ± 10.2 nM. The association rate constant (k_on_) was determined to be 7.11 ± 0.11 × 10^3^ M^-1^s^-1^. Notably, the dissociation rate constant (k_off_) of candidalysin from heparin was quite low at 2.30 ± 0.14 × 10^-4^ (×10^-5^) s^-1^, indicating that the half-life of the heparin-candidalysin complex likely exceeds 50 minutes. Therefore, candidalysin binds to heparin with high affinity and forms a stable complex.

### Exogenous sulfated GAGs protect epithelial cells from candidalysin-induced damage

All naturally occurring GAGs are negatively charged linear polysaccharides ^28–30^. The finding that hyaluronic acid failed to protect epithelial cells from candidalysin-induced damage suggested that protection is due to biochemical, conformational or other stereochemical factors beyond positively charged candidalysin molecules binding to negatively charged GAGs. To investigate the specificity of GAGs in reducing epithelial cell damage by candidalysin, we tested the protective effects of the highly carboxylated, negatively charged polysaccharide alginate. Up to the highest concentration tested (128 μg/ml), alginate did not ameliorate candidalysin-induced damage (Supplementary Fig. 4a). This result confirmed that the candidalysin interaction with some GAGs cannot be explained by charge-charge interactions alone. Major differences among hyaluronic acid, alginate and other naturally occurring GAGs, such as heparin, heparan sulfate and chondroitin sulfate, are the extent and stereochemical positioning of sulfation (Supplementary Fig. 3a). To determine whether sulfation of GAGs is required for their protective effects, we analyzed the sulfated and non-sulfated GAG analogs, dextran sulfate and dextran (see structure in Supplementary Fig. 3b). We found that dextran sulfate protected epithelial cells from candidalysin-induced damage in a dose-dependent manner, whereas non-sulfated dextran had no effect (Fig. 2d). Dextran sulfate with molecular masses ranging from 10 kDa to 500 kDa were similarly protective, indicating that the protective effect was independent of molecular mass (Supplementary Fig. 4b).

Heparan sulfate and heparin (a highly sulfated form of heparan sulfate) have a linear structure, whereas dextran sulfate has a branched structure. Yet, all of these molecules protected cells against candidalysin-induced damage. These results suggest that the protective effects of GAGs and GAG analogs may be largely independent of the structure of the backbone molecule to which the sulfate moiety is attached. In support of this hypothesis, we found that the cyclic hexasaccharide alpha-cyclodextrin (see structure in Supplementary Fig. 3b) was also protective, but only when it was sulfated (Fig. 2d). These sulfated GAGs and their analogs were also protective against 70 μM candidalysin, a highly lethal concentration, although the extent of protection was decreased (Supplementary Fig. 4c). The protection effects of dextran sulfate were maintained for at least 24 h (Supplementary Fig. 4d). These data suggest that sulfation of GAG or its analogs is a critical factor for protection from candidalysin-induced cell damage, whereas the molecular mass or the structure of the backbone molecule is less important.

To determine if the location of the sulfate moiety on the backbone molecule also affected binding to candidalysin, we used SPR to test the capacity of various de-sulfated heparins to compete with native heparin for binding to candidalysin. While pre-incubation of candidalysin with heparin and 2-*O*-desulfated heparin inhibited candidalysin binding to chip-immobilized heparin, *N-*desulfated heparin and 6-*O*-desulfated heparin failed to inhibit candidalysin binding to heparin (Fig. 2e-g). Thus, both the *N*-sulfo and 6-*O*-sulfo groups on heparin are required for candidalysin binding, underscoring the importance of stereospecific sulfate modifications on GAGs during their interaction with candidalysin.

Using an SPR-based competition assay, we also assessed the capacity of various GAGs to inhibit the interaction between candidalysin and chip-immobilized heparin. Dextran, chondroitin sulfate A (chondroitin-4-sulfate), chondroitin sulfate B (dermatan sulfate), chondroitin sulfate C (chondroitin-6-sulfate), and heparan sulfate did not significantly inhibit candidalysin binding to chip-immobilized heparin (Supplementary Fig. 4e, f). By contrast, heparin and dextran sulfate both inhibited binding, with dextran sulfate reducing binding by >95% (Supplementary Fig. 4e, f). These results suggest that dextran sulfate has the highest binding affinity for candidalysin, followed by heparin and then the other GAGs that were tested.

### GAGs facilitate polymerization and/or aggregation of candidalysin

It has been previously demonstrated that candidalysin is able to form linear and circular structures (loops) in aqueous solution prior to inserting into the cell membrane and forming pores ^4^. By transmission electron microscopy, we observed that candidalysin formed the expected linear polymers and loops when added to a solid substrate in the presence of the vehicle control or dextran (Supplementary Fig. 5a). The loops, which are expected to become pores when inserted into membrane, were replaced by aggregates in the presence of dextran sulfate or heparan sulfate. Atomic force microscopy (AFM) imaging of candidalysin on a solid substrate also showed that free dextran sulfate and heparan sulfate caused significant aggregation of candidalysin (Supplementary Fig. 5a). These topographically high aggregates appeared as bright white structures under AFM. Dextran, on the other hand, displayed minimum effect on candidalysin structures. Using AFM imaging of candidalysin on a 1,2-dioleoyl-sn-glycero-3-phosphocholine (DOPC) bilayer, we observed that candidalysin formed pores when added to the bilayer either alone or in the presence of dextran (Fig. 3a). These pores were absent when candidalysin was mixed with either heparan sulfate or dextran sulfate. Take together, these data suggest that binding to free sulfated GAGs leads to polymerization (aggregation) of candidalysin. This interaction with sulfated GAGs that are expressed on the cell surface likely increases the concentration of the peptide on the cell membrane to promote its pore-forming ability. On the other hand, the aggregates observed in the presence of free GAGs are probably non-functional, because they reduce the number of pore-competent loops.

**Fig. 3.**
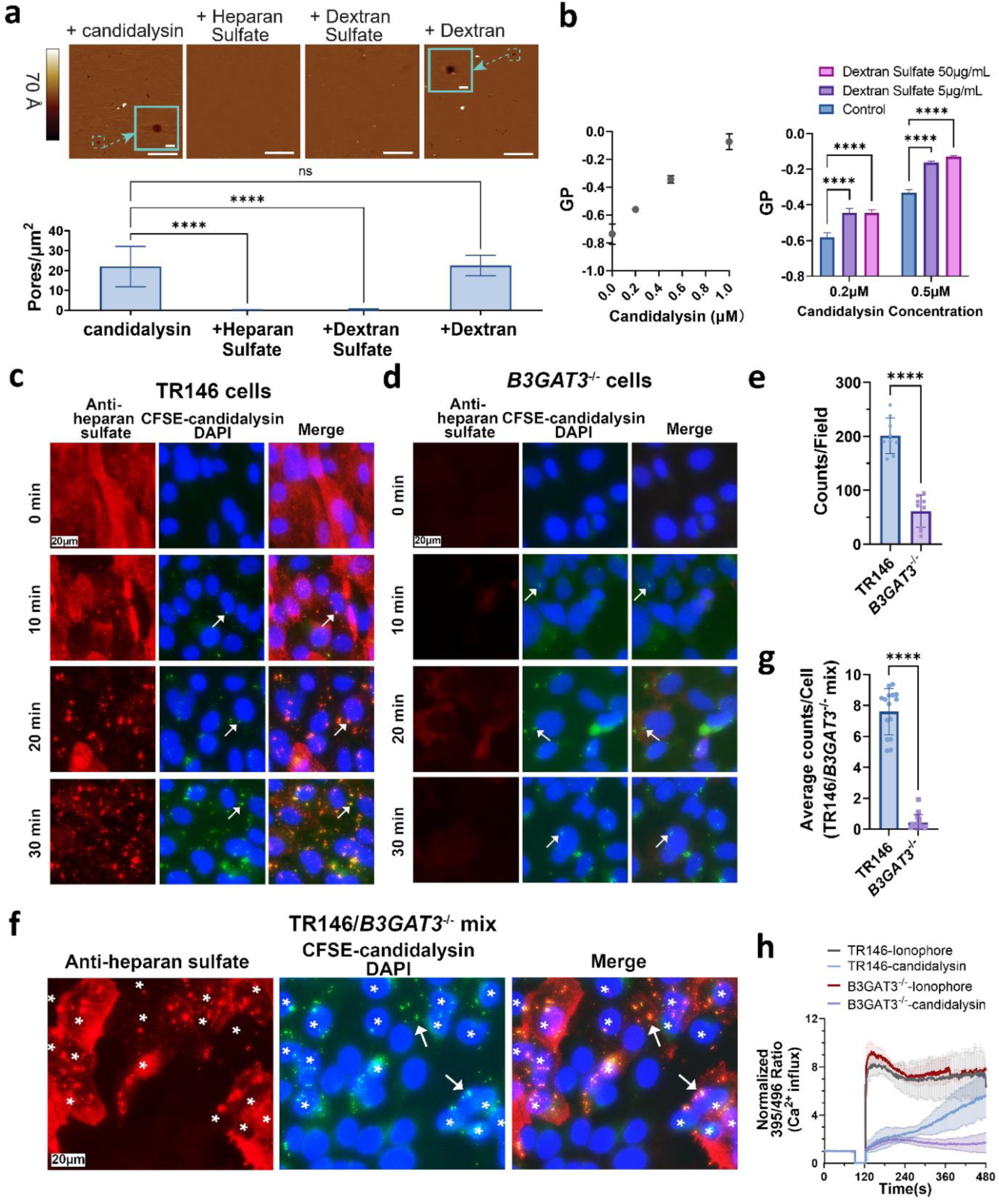
GAGs enhance candidalysin polymerization/aggregation. **a,** Atomic force microscopy images of supported 1,2-dioleoyl-sn-glycero-3-phosphocholine (DOPC) lipid bilayers exposed to candidalysin (333 nM) in the presence and absence of 5 μg/ml dextran, dextran sulfate and heparan sulfate (upper panel), and the quantification of the number of pores per μm^2^ for each treatment (lower panel). Scale bar: 200 nm. Results are mean ± SD of 3 experiments. **b,** Effects of candidalysin concentration on the general polarization (GP) score using a membrane-less C-laurdan assay (left panel). Data were obtained after incubating the indicated concentrations of candidalysin in 1 µM C-laurdan for 4 h. The effects of the indicated concentration of dextran sulfate on the GP scores of 0.2 μM and 0.5 μM candidalysin (right panel). Results are mean ± SD of 3 independent experiments. **c,** Co-localization of candidalysin with heparan sulfate on the surface of wild-type TR146 oral epithelial cells incubated with CSFE-labeled candidalysin (10 μM) for the indicated times. Arrows indicate representative candidalysin aggregates. Scale bar: 20 μm. **d,** Interactions of CFSE-labeled candidalysin (10 μM) with a GAG-deficient *B3GAT3*^-/-^ oral epithelial cell line for the indicated times. Arrows indicate representative candidalysin aggregates. Scale bar: 20 μm. **e**, Quantification of candidalysin aggregates per microscopic field on TR146 wild-type and *B3GAT3*^-/-^ epithelial cells after exposure to CFSE-labeled candidalysin for 20 min. Results are mean ± SD of 3 random microscopic fields per experiment from 3 independent experiments. **f,** Interactions of CFSE-labeled candidalysin (10 μM) with a 1:1 mixture of wild-type and GAG-deficient *B3GAT3*^-/-^ oral epithelial cells. Arrows indicate representative candidalysin aggregates. The wild-type cells stain for heparan sulfate (red) and are marked with asterisks. Scale bar: 20 μm. **g,** Quantification of candidalysin aggregates per TR146 or *B3GAT3*^-/-^ cell in the mixed population of TR146/*B3GAT3*^-/-^ cells after exposure to CFSE-labeled candidalysin (10 μM) for 20 min. Results are mean ± SD of 5 random fields per experiment from 3 independent experiments. **h**, Normalized fluorescence intensity of Ca^2+^-bound Indo-1/AM dye in TR146 cells and *B3GAT3*^-/-^ cells treated with calcium ionophore A23187 (1 μM) or candidalysin (10 μM) as determined by flow cytometry. 0-90s represents the baseline intensity and data collection was resumed at 120s after the reagent was added to cells. Results are mean ± SD of 4 independent experiments. The p values for candidalysin treated TR146vs*B3GAT3*^-/-^ were ns from 1-317s, * from 318-337s, ** from 338-365s, and *** from 366-481s. ns, not significant; *, p<0.05; **, p<0.01; ***, p<0.001; ****, p<0.0001 by one-way ANOVA with the Dunnett’s multiple comparisons test (**a, b**), by the unpaired, two-sided Student’s t test (**e, g**) and by Two-way ANOVA with Tukey’s multiple comparisons tests (**h**).

To test this hypothesis, we measured the response of 6-dodecanoyl-2-[N-methyl-N- (carboxymethyl)amino]naphthalene (C-laurdan) to candidalysin. C-laurdan has spectral sensitivity to the polarity of its environment, reflecting local water content. When C-laurdan is dissolved in a polar solvent, its emission spectrum shifts by approximately 50 nm ^31,32^. We analyzed the effects of various concentrations of candidalysin on the fluorescence emission of C-laurdan. A generalized polarization (GP) score of C-laurdan was calculated as described in the Methods. The GP score increases as binding to or polymerization/aggregation of candidalysin reduces the local hydration of the dye (Supplementary Fig. 5b). We found that the GP score rose as the concentration of candidalysin increased, which promotes peptide polymerization (Fig. 3b). Incubation of dextran sulfate with candidalysin further increased the GP score, suggesting greater candidalysin polymerization and/or aggregation (Fig. 3b). In the absence of candidalysin, dextran sulfate had no effect on the GP score (Supplementary Fig. 5c). We also found that candidalysin was able to cause membrane permeation when added to artificial vesicles composed of DOPC or 1-palmitoyl-2-oleoyl-sn-glycero-3-phosphocholine (POPC), which was delayed when free dextran sulfate was added to the medium before candidalysin (Supplementary Fig. 5d). Thus, our data suggest that dextran sulfate enhances the self-assembly of candidalysin in an aqueous solution and delays pore formation.

Next, we investigated whether GAGs on the epithelial cell surface altered the interactions of candidalysin with the cells. Candidalysin was fluorescently labeled with carboxyfluorescein succinimidyl ester (CFSE) and added to TR146 oral epithelial cells. The CFSE-labeled candidalysin remained functional, as it damaged the epithelial cells similarly to unlabeled candidalysin (Supplementary Fig. 5e). At various time points, the treated cells were then fixed with paraformaldehyde and the heparan sulfate on the cell surface was fluorescently labeled with an anti-heparan sulfate antibody for imaging by indirect immunofluorescence microscopy. When added to wild-type epithelial cells, the CFSE-labeled candidalysin formed aggregates that increased in number and size over time (Fig. 3c). These aggregates colocalized with heparan sulfate (Supplementary Fig. 5f), suggesting that candidalysin indeed interacts with cell surface GAGs. Intriguingly, candidalysin caused a progressive loss of surface exposed heparan sulfate, so that by 30 min very little heparan sulfate could be detected. Taken together, our results indicates that candidalysin interacts with GAGs on the epithelial cell surface.

To determine how the presence of GAGs influences the interactions of candidalysin with oral epithelial cells, we repeated this experiment using *B3GAT3*^-/-^ cells, which lack cell surface GAGs (Supplementary Fig. 2c-d) and are more resistant to damage from candidalysin relative to wild-type cells (Fig. 1b). We observed that candidalysin still formed aggregates on the cell surface of the mutant cells, but these aggregates were less numerous and more diffuse than those that formed on wild-type cells (Fig. 3c-e). These results are consistent with the model that the reduced candidalysin aggregation on *B3GAT3*^-/-^ cells is due to the absence of cell surface GAGs.

We also added CSFE-candidalysin to a mixed population of wild-type and GAG deficient *B3GAT3*^-/-^ cells to mimic the conditions under which the CRISPR library was screened. Candidalysin preferentially localized to the GAG sufficient wild-type cells (Fig. 3f-g). This preferential binding to GAG-expressing cells likely protected the GAG deficient cells from candidalysin-induced damage, providing an explanation for why genes required for GAG biosynthesis were among the top hits in our CRISPR screen. These data further support the idea that sulfated GAGs bind to candidalysin and facilitate its polymerization on the cell surface.

The polymerization of candidalysin results in the formation of pores in the plasma membrane, leading to an influx of calcium into the cell^1^. To determine if the observed reduction in candidalysin aggregates in the *B3GAT3*^-/-^ cells resulted in decreased pore formation, we used flow cytometry to measure the levels of intracellular calcium in epithelial cells after treatment with candidalysin. We found that in response to candidalysin, the levels of intracellular calcium rose more slowly in the GAG-deficient *B3GAT3*^-/-^ cells as compared to wild-type TR146 cells (Fig. 3h). By contrast, the kinetics of intracellular calcium increase into these cells were similar when they were incubated with the A23187 calcium ionophore. These results indicate that formation of functional candidalysin pores occurs more slowly in the absence of cell surface GAGs.

### GAGs are required for maximal epithelial cell invasion, damage, and proinflammatory response by live *C. albicans*

To test if sulfated GAGs are required for *C. albicans* pathogenicity, we analyzed the interactions of live *C. albicans* with epithelial cells *in vitro*. We found that relative to wild-type TR146 cells, the GAG-deficient *B3GAT3*^-/-^, *B3GALT6*^-/-^, and *XYLT2*^-/-^ epithelial cells were less susceptible to damage caused by live *C. albicans* (Fig. 4a). Although live *C. albicans* cells had slightly reduced cell-association (a measure of adherence) to the GAG-deficient cells, their endocytosis by these cells was substantially decreased (Fig. 4b-c). Thus, epithelial cell GAGs are required for maximal *C. albicans* adherence, invasion, and damage *in vitro*.

**Fig. 4.**
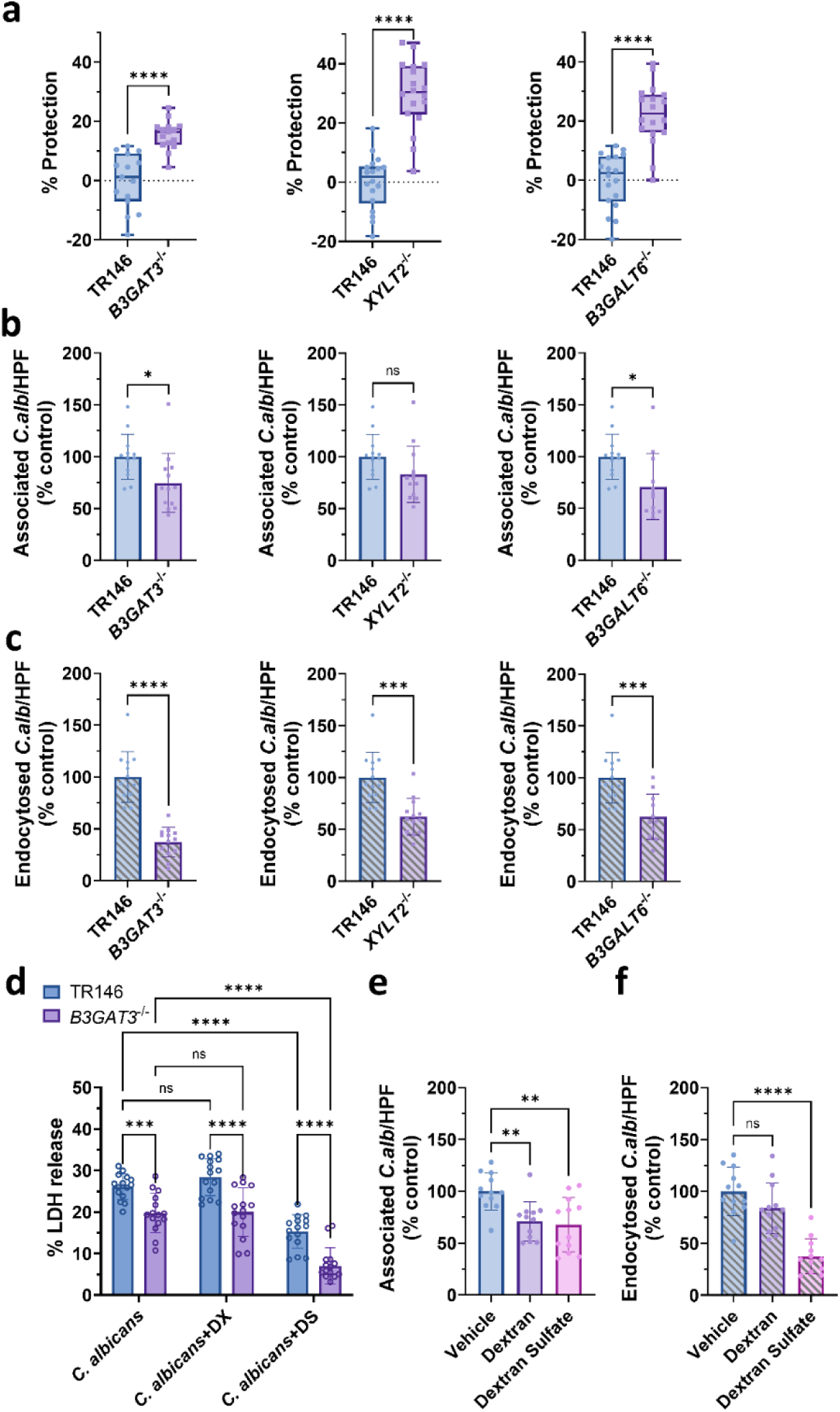
Sulfated GAGs mediate epithelial cell invasion, damage, and stimulation of oral epithelial cells by live *C. albicans*. **a,** Protection against damage caused by 5 h of infection with live wild-type *C. albicans* at a multiplicity of infection (MOI) of 5 in *B3GAT3*^-/-^, *XYLT2*^-/-^, and *B3GALT6*^-/-^ mutant cells relative to wild-type TR146 epithelial cells, as measured by an XTT assay. **b,c,** *C. albicans* association with (**b**) and endocytosis by (**c**) wild-type and *B3GAT3*^-/-^, *XYLT2*^-/-^, and *B3GALT6*^-/-^ mutant epithelial cells. The average number of organisms per high-power field that were associated with and endocytosed by TR146 cells were 12.84 ± 4.97 and 2.91 ± 1.27, respectively. **d,** Effects of dextran and dextran sulfate (100 μg/ml) on damage to TR146 and *B3GAT3*^-/-^ cells caused by live *C. albicans* SC5314 at an MOI of 5 for 5 h, as measured by an LDH release assay. **e, f,** Effects of dextran and dextran sulfate (100 μg/ml) on wild-type *C. albicans* association with (**e**) and endocytosis by (**f**) the indicated epithelial cells. The average number of organisms per high-power field that were associated with and endocytosed by TR146 cells were 9.64 ± 2.68 and 1.88 ± 0.56, respectively. Results are the mean ± SD of three experiments, each performed in triplicate. ns, not significant; *, p<0.05; **, p<0.01; ***, p<0.001; ****, p<0.0001 by unpaired, two-tailed Student’s test (**a-c**) or one-way ANOVA with Dunnett’s multiple comparisons test (**d-f**).

The finding that dextran sulfate blocked candidalysin-induced damage to epithelial cells suggested that it would also protect epithelial cells from damage caused by live *C. albicans.* To investigate this possibility, we assessed the extent of damage caused by *C. albicans* to TR146 epithelial cells in the presence or absence of dextran sulfate. To facilitate comparison with previous studies ^1,3,33,34^, we measured the extent of epithelial cell damage using an LDH release assay. As shown in Figs. 4a and c, this assay yielded similar results to the XTT assay. We found that dextran sulfate partially protected the epithelial cells from damage caused by wild-type *C. albicans* SC5314, whereas dextran had no effect (Fig. 4d). Although *C. albicans* caused less damage to *B3GAT3*^-/-^ cells than to wild-type TR146 cells, dextran sulfate caused a further reduction in damage (Fig. 4d). Because the candidalysin-deficient *ece1*Δ/Δ strain did not cause detectable damage to TR146 or *B3GAT3*^-/-^ cells (Supplementary Fig. 5g), we were unable to determine the effects of dextran sulfate on epithelial cell damage caused by this strain. These data indicate that dextran sulfate reduces *C. albicans*-induced epithelial cell damage by both GAG-dependent and -independent mechanisms.

Next, we assessed the effects of dextran and dextran sulfate on *C. albicans* adherence to and endocytosis by oral epithelial cells, testing both the wild-type and *ece1*Δ/Δ mutant strains. Both dextran and dextran sulfate caused a modest, yet statistically significant reduction in the adherence of wild-type *C. albicans* to oral epithelial cells (Fig. 4e). Both compounds trended towards reducing adherence of the *ece1*Δ/Δ mutant, but this decrease was not significant (Supplementary Fig. 5h). Dextran sulfate decreased the endocytosis of both the wild-type strain and the *ece1*Δ/Δ mutant by approximately 60% (Fig. 4f, Supplementary Fig. 5i). Collectively, these results indicate that dextran sulfate can not only decrease the extent *C. albicans*-induced damage to oral epithelial cells, but also the capacity of the organism to adhere to and invade these cells. As candidalysin does not mediate adherence to or invasion of oral epithelial cells, the inhibitory effects of dextran sulfate on these processes is independent of its effects on candidalysin.

Infection of epithelial cells with some strains of *C. albicans*, such as SC5314, induces a robust pro-inflammatory response, and candidalysin has been identified as one of the main inducers of this response ^3,12^. Exposure of oral epithelial cells to live C*. albicans* or candidalysin triggers the phosphorylation of the epidermal growth factor receptor (EGFR) ^35^, ERK1/2 signaling cascade, and consequently, phosphorylation of the c-Fos transcription factor ^12,14^, a result we confirmed (Fig. 5a). We found that dextran sulfate, but not non-sulfated dextran, inhibited the phosphorylation of EGFR, ERK1/2, and c-Fos induced by either wild-type *C. albicans* or candidalysin in both GAG-sufficient TR146 and GAG-deficient *B3GAT3*^-/-^ oral epithelial cells (Fig. 5a, Supplementary Fig. 6a). As expected, the *ece1*Δ/Δ mutant did not stimulate phosphorylation of EGFR, ERK1/2, and c-Fos above basal levels in wild-type and *B3GAT3*^-/-^ oral epithelial cells. These results indicate that while GAGs are not required for *C. albicans* or candidalysin to stimulate oral epithelial cells, dextran sulfate can inhibit the response to both stimuli.

**Fig. 5.**
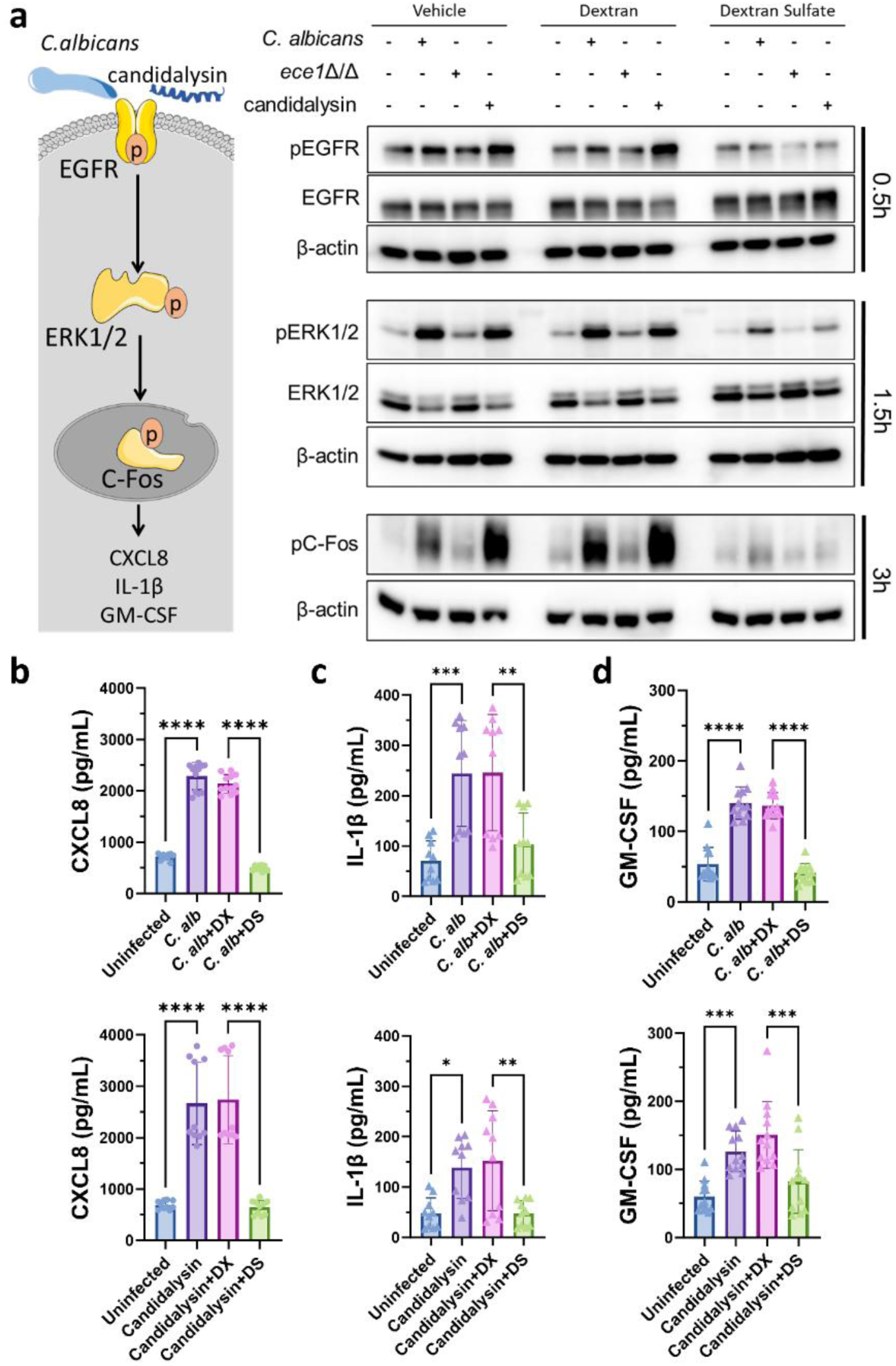
Sulfated GAGs mediate stimulation of oral epithelial cells by candidalysin and live *C. albicans*. **a,** Simplified model of epithelial immune signaling in response to *C. albicans* and candidalysin (left panel). Western blot analysis of the effects of dextran (DX, 100 μg/ml) and dextran sulfate (DS, 100 μg/ml) on the phosphorylation of epidermal growth factor receptor (EGFR), extracellular regulated kinase1/2 (ERK1/2) and the C-Fos transcription factor in TR146 cells induced by wild-type *C. albicans* SC5314 or the *ece1*Δ/Δ mutant at MOI of 5, or candidalysin (10 μM) at the indicated time points (right panel). Shown are representative results of 3 independent experiments. **b-d,** Effects of dextran and dextran sulfate (100 μg/ml) on the production of CXCL8 (**b**), IL-1β (**c**) and GM-CSF (**d**) by TR146 cells infected with *C. albicans* SC5314 (top, MOI=5) or incubated with candidalysin (bottom, 10 μM) for 6 h. Results in (**b-d)** are mean ± SD of 3 experiments, each performed triplicate. **, p < 0.01; ****, p < 0.0001 by one-way ANOVA with Dunnett’s multiple comparisons test (**b-d**).

Activation of oral epithelial cells by either live *C. albicans* or candidalysin induces a pro-inflammatory transcriptional response, resulting in the secretion of cytokines, such as CXCL8, IL-1β, and GM-CSF (Fig. 5b-d). Dextran sulfate, but not dextran significantly decreased the production of these cytokines in wild-type TR146 cells. (Fig. 5b-d). Interestingly, while exposure of *B3GAT3*^-/-^ oral epithelial cells to live, wild-type *C. albicans* stimulated the production of all three of these cytokines, candidalysin only induced the secretion of CXCL8 and GM-CSF but not IL-1β (Supplementary Fig. 6b). These responses were almost completely inhibited by dextran sulfate, but not dextran. The *ece1*Δ/Δ mutant did not induce a detectable increase in secretion of any of these cytokines by either *B3GAT3*^-/-^ or wild-type TR146 cells (Supplementary Fig. 6b-c). Thus, dextran sulfate interferes with the signaling cascade that induces the secretion of pro-inflammatory mediators in response to *C. albicans* and candidalysin.

### Dextran sulfate treatment ameliorates vulvovaginal candidiasis

The body of evidence suggested that dextran sulfate could interfere with *Candida* pathogenicity *in vivo.* To evaluate this hypothesis, we tested the capacity of dextran sulfate to protect epithelial cells from *C. albicans* in a mouse model of mucosal candidiasis. We could not test the effects of dextran sulfate in the mouse model of oropharyngeal candidiasis because the compound would need to be administered continuously to prevent it from being cleared from the oropharynx by salivary flow. To circumvent this problem, we tested the efficacy of dextran sulfate in the mouse model of vulvovaginal candidiasis, where it remained in the vagina for an extended period of time. Prior to the mouse studies, we verified that dextran sulfate protected the A431 vaginal epithelial cell line from damage caused by candidalysin (Fig. 6a). Dextran sulfate also diminished the production of IL-1β and CXCL8 by these cells induced by either live *C. albicans* or candidalysin (Fig. 6b).

**Fig. 6.**
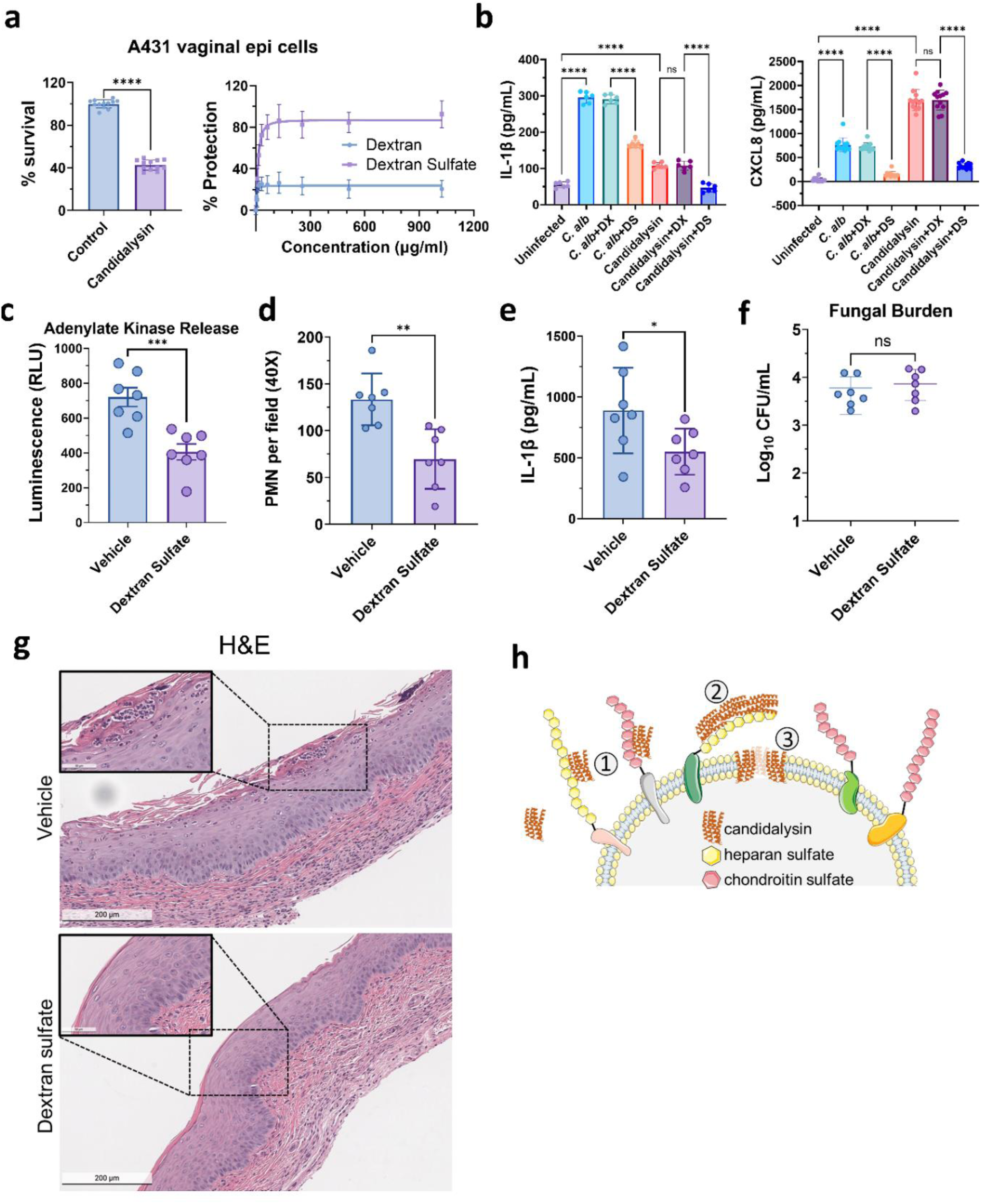
Dextran sulfate protects vaginal epithelial cells from damage and inhibits pro-inflammatory cytokine production *in vitro* and during murine vulvovaginal candidiasis. **a,** Survival of A431 vaginal epithelial cells after incubation with 30 μM candidalysin for 6 h (left). Protection from candidalysin-induced damage by dextran and dextran sulfate (right). **b,** Effects of dextran and dextran sulfate (100 μg/ml) on the production of IL-1β and CXCL8 by vaginal epithelial cells incubated with *C. albicans* (MOI=5) or candidalysin (10 µM) for 24 h. Results in (**a, b**) are mean ± SD of 3 experiments, each performed in triplicate. **c-g,** C57BL/6 mice were administered either dextran sulfate or vehicle alone intravaginally prior to intravaginal inoculation with *C. albicans* and once daily thereafter. After 3 days of infection, the concentration of adenylate kinase (a measure of host cell damage) (**c**), neutrophils (PMN) (**d**), IL-1β (**e**), and fungal colony forming units (CFU) (**f**) in the vaginal lavage fluid was determined. Results in **c-f** are the mean ± SD 2 experiments, one that used 3 mice per condition and the other that used 4 mice per condition. **g**, Representative images of hematoxylin and eosin (H&E) stained histological sections of vaginal tissue day3 post-infection. The scale bar represents 200 µm in full images and 50 µm in magnified images in the lower right. **h**, A proposed working model of how cell surface GAGs facilitate candidalysin polymerization and pore-formation into host cells. Free candidalysin can bind to GAGs such as heparan sulfate or chondroitin sulfate that are attached to cell surface glycoproteins (1). Binding to these cell surface GAGs enhances the aggregation and polymerization of candidalysin (2), which facilitates the formation of candidalysin pores that insert into the host cell plasma membrane (3), leading to host cell damage. ns, not significant; *, *p* < 0.05; **, *p* < 0.01; ***, *p* < 0.001; ****, *p* < 0.0001 by unpaired, two-sided Student’s t-test (**a, c-f**) or one-way ANOVA with Dunnett’s multiple comparisons test (**b**).

In the C57BL/6 mouse model of vulvovaginal candidiasis, daily administration of dextran sulfate intravaginally significantly decreased the extent of host cell damage in the vagina (Fig. 6c). Dextran sulfate also reduced vaginal inflammation, as reflected by reduced levels of neutrophils and IL-1β (Fig. 6d-e). By contrast, this treatment did not significantly alter the vaginal fungal burden (Fig. 6f). Histopathologic analyses verified that the dextran sulfate-treated mice exhibited less damage to the vaginal epithelium and lower neutrophil accumulation compared to mice treated with vehicle alone (Fig. 6g). We also tested the effects of dextran sulfate on the outcome of vaginal candidiasis in CD-1 mice, which are reported to be resistant to this infection ^35^. In the mice that achieved a high vaginal fungal burden, dextran sulfate treatment significantly reduced tissue damage and decreased neutrophil accumulation (Supplementary Fig. 7a and b). While there was a trend towards reduced IL-1β in the dextran sulfate-treated mice, this difference was not statistically significant (*p* = 0.08) (Supplementary Fig. 7c). Similar to what was found with C57BL/6 mice, dextran sulfate treatment had no effect on the vaginal fungal burden (Supplementary Fig. 7d). Collectively, these data indicate that dextran sulfate ameliorates vaginal candidiasis in two different mouse strains.

## Discussion

Through a genome-wide CRISPR knockout screen, we identified three genes, *B3GAT3*, *B3GALT6*, and *XYLT2*, that are required for maximal susceptibility to candidalysin-induced host cell damage. The products of each of these genes catalyze linear early steps in GAG biosynthesis in host cells. We subsequently found that candidalysin binds to sulfated GAGs *in vitro* and that the GAG analog dextran sulfate ameliorates epithelial cell damage and reduces cytokine release induced by both candidalysin and live *C. albicans.* The protective effects of sulfated dextran were demonstrated *in vitro* and in the mouse model of vulvovaginal candidiasis. Collectively, these data indicate that GAGs play a key role in the interactions of candidalysin and *C. albicans* with host epithelial cells.

It has recently been found that glycans are critical host receptors for bacterial and viral virulence factors ^22,36–40^. Even so, the role of host glycans is poorly understood with respect to fungal virulence factors. Although previous studies indicated that candidalysin interacts with cell membranes ^4,10^, the current work shows that a more specific interaction with GAGs is also needed for the full activity of this toxin. Thus, the work presented here further extends the observation of the importance of glycan receptors to a fungal virulence factor.

The three genes in the GAG biosynthesis pathway, *B3GAT3*, *B3GALT6*, and *XYLT2,* that we discovered were required for maximal candidalysin-induced damage are among the top hits in numerous other CRISPR screens aimed at identifying key host factors for viral and bacterial pathogenicity ^36,37,39–42^. A notable difference is that many of the other screens, but not the one performed here, also identified *EXT1*, *EXT2*, and *EXT3*, which encode glycosyltransferases that are crucial for linear chain extension during the synthesis of heparan sulfate ^36,37,43^. While these enzymes are only required for the biosynthesis of heparan sulfate, B3gat3, B3galt6 and Xylt2 are essential for the early steps in the biosynthesis of all natural occurring sulfated GAGs, including heparan sulfate, chondroitin sulfate and dermatan sulfate. Thus, our results indicate that candidalysin can interact with most sulfated GAGs to enhance host cell damage. In support of this conclusion, we demonstrated that exogenous heparin, heparan sulfate and chondroitin sulfate could protect epithelial cells against candidalysin-induced damage in a competition assay.

Our screen also identified *SLC39A9* as a gene required for maximal candidalysin-induced damage. This gene was previously identified in a CRISPR screen for resistance to the subtilase cytotoxin SubAB ^22^. Interestingly, deletion of *SLC39A9* was shown to reduce N- and O-glycan modification of cell surface proteins. Thus, it is possible that SLC39A9 is required for the surface expression of GAGs that bind to candidalysin.

Recently, the intracellular host cell ligands of various Ece1 peptides, including candidalysin, were identified using a yeast two-hybrid screen ^44^. The results of this screen indicated that candidalysin binds to CCNH, which is a regulatory subunit of the CDK-activating kinase complex. This binding inhibits DNA repair, both *in vitro* and in the mouse model of oropharyngeal candidiasis. As expected, the yeast two hybrid screen did not identify B3gat3, B3galt6, or Xylt2 because our data indicate that candidalysin binds to GAGs, which are the products of these enzymes, and not the enzymes themselves.

Our exploration into the impact of GAGs on candidalysin polymerization and aggregation through AFM and TEM suggests that GAGs facilitate candidalysin polymerization or aggregation. Our immunofluorescence assays further confirmed that candidalysin interacts with GAGs on the epithelial cell surface, and GAG deficient cells demonstrate less efficient candidalysin attachment, aggregation and pore-formation. This body of evidence strongly supports our hypothesis that linear GAGs provide a platform for candidalysin to form aggregates on the cell surface and enhance the cytotoxic functionality of candidalysin (Fig. 6h).

Exogenous sulfated GAGs and dextran sulfate almost completely blocked epithelial cell damage by competitive binding to candidalysin. However, the GAG-deficient *B3GAT3*^-/-^, *B3GALT6*^-/-^ and *XYLT2*^-/-^ mutant epithelial cells were only partially protected from candidalysin-induced damage. This result is concordant with previous findings ^4^ and our own data, which demonstrate that candidalysin can induce membrane damage in GAG-free liposomes composed of the synthetic lipids POPC or DOPC. Taken together, our results indicate that while GAGs enhance the polymerization of candidalysin at or near the cell surface, this process can occur in the absence of cell surface GAGs, albeit less efficiently.

Candidalysin displays a net positive charge at physiological pH due to the three lysine residues in its C-terminus ^1,45^. Several publications have demonstrated that the cytotoxicity of candidalysin can be mitigated by negatively charged molecules such as albumin, heparin, and polyacrylic acid ^45–47^. In contrast, we found that the negatively charged hyaluronic acid and alginitic acid failed to protect cells from candidalysin-induced damage. This seeming divergence in results could be due to the fact that the concentrations of hyaluronic acid and alginitic acid used in our experiments were 1- to 2-log lower than the concentrations of albumin and polyacrylic acid used in the other studies. Of note, our surface plasmon resonance assays revealed a strong binding affinity between candidalysin and heparin (K_D_ = 3.24 × 10^-8^ M), highlighting the specificity and strength of this interaction. Furthermore, the importance of *N*- and 6-*O*-sulfate groups on heparin for the heparin-candidalysin interaction implies that charge density and/or stereospecific sulfation patterns, rather than electrostatics, structural attributes or molecular weight alone determine the interaction strength of GAGs with candidalysin.

The data presented here also indicate that cell surface GAGs are required for maximal *C. albicans* adherence to and invasion of oral epithelial cells. Deletion of *B3GAT*^-^, *B3GALT6*, or *XYLT2* modestly reduced adherence and substantially reduced invasion of *C. albicans* into these cells. Dextran sulfate also inhibited epithelial cell invasion in a candidalysin-independent manner. The likely explanation for these results is that GAGs are required for the proper function of the epithelial cell receptors that mediate the adherence and endocytosis of *C. albicans*. For example, it has been reported that glycosylation is necessary for the proper function of both EGFR and E-cadherin, which are two key epithelial cell receptors for *C. albicans*. Thus, it is likely that the protective effects of dextran sulfate, both *in vitro* and *in vivo* were due to both neutralization of candidalysin and inhibition of epithelial cell receptor function.

We found that dextran sulfate not only protected epithelial cells from *C. albicans-*induced damage, but also diminished signaling through EGFR, ERK1/2 and c-Fos, leading to reduced release of proinflammatory cytokines. As these pathways are known to be activated by candidalysin, it is probable that this inhibition was due in part to neutralization of candidalysin by dextran sulfate. It has also been reported that heparin-binding epidermal growth factor-like (HB-EGF) growth factor activates EGFR in a paracrine manner and that this activation can be blocked by heparin ^48^. Thus, it is possible that dextran sulfate also inhibited epithelial cell stimulation by directly blocking EGFR activation, similarly to heparin.

A central finding was that in the mouse model of vulvovaginal candidiasis, dextran sulfate not only reduced host cell damage, but also decreased cytokine levels and neutrophil accumulation at the infection site. Notably, dextran sulfate had no effect on vaginal fungal burden. This pattern of reduced vaginal damage and inflammation, but no change in fungal burden is very similar to what has been seen in mice infected with the candidalysin-deficient *ece1*Δ/Δ mutant. This similarity provides additional evidence that the salutary effects of dextran sulfate were due at least in part to its neutralization of candidalysin. However, it is also likely that dextran sulfate inhibited the pathogenic interactions of *C. albicans* with vaginal epithelial cells by additional mechanisms, such as inhibiting *C. albicans* invasion.

In summary, our study elucidates the intricate relationship between candidalysin and cell surface GAGs, uncovering specific host factors that are required for maximal candidalysin-induced damage. The protective effects of exogenous GAGs, particularly dextran sulfate, present novel therapeutic avenues for mitigating the impact of *C. albicans* infection. These findings contribute to our understanding of the molecular mechanisms underlying candidalysin-induced damage and open new possibilities for targeted antifungal interventions.

## Materials and Methods

### Fungal strains

*C. albicans* SC5314 was used as the wild-type strain. The candidalysin-deficient *ece1*Δ/Δ mutant was constructed as described ^49^. The organisms were grown overnight in yeast extract, peptone, dextrose (YPD) broth at 30°C in a shaking incubator. The next day, the yeast-phase organisms were harvested by centrifugation, washed twice with PBS, enumerated with hemacytometer and then resuspended in tissue culture medium.

### Candidalysin

Candidalysin (NH2-SIIGIIMGILGNIPQVIQIIMSIVKAFKGNK-COOH) was produced by solid-phase peptide synthesis using F-MOC chemistry (BioMatik). It was purified to >95% purity by reverse-phased high-performance liquid chromatography. Authentication of sequence and molecular mass (3311.7 daltons [g/mol]) were achieved using liquid chromatography mass spectroscopy. Stock solutions of 10 mM candidalysin were prepared in pure water and stored at −80°C.

To label candidalysin with carboxyfluorescein succinimidyl ester (CFSE), 10 μL of 10 mM candidalysin in water was mixed with 90 μL DMSO to create a 1 mM candidalysin stock. Next, CFSE (Biolegend, #423801) was added to the 1 mM candidalysin stock at a final concentration of 5 μM. The mixture was vortexed and then incubated in the dark at room temperature for 20 min. After the labeled candidalysin was prepared, 10 μL was mixed with 1 mL of serum-free, supplement free medium and then incubated with TR146 cells on glass coverslips in a 24-well plate for various times. The cells were then fixed with paraformaldehyde before subsequent immunofluorescence staining.

### Cell culture

The TR146 human buccal epithelial squamous cell carcinoma cells (MilliporeSigma) ^50^ were incubated in DMEM/F12 medium (Gibco, #11320033) with 10% fetal bovine serum and 1x penicillin-streptomycin (Gibco, #15140122). A431 vulval epithelial cells (MilliporeSigma) and HEK-293T cells (ATCC) were cultured in DMEM media supplemented with 10% heat-inactivated fetal bovine serum and 2 mM glutamine (Corning, #25005-CL). The cells were switch to serum free medium the day before the experiments.

### Antibodies and reagents

Mouse anti-Cas9 (#sc517386), mouse anti-hGBF1 (#sc136240), mouse anti-c-Fos (#sc8047), mouse anti-hB3GAT3 (#sc390526), mouse anti-hXYLT2 (#sc374134) antibodies were purchased from Santa Cruz Biotechnology, and mouse anti-Heparan Sulfate antibody (clone F58-10E4, #3702551) was ordered from AMSBIO. Rabbit anti-phospho-c-Fos (Ser32, #5348), rabbit anti-phospho-EGFR (Tyr1068, #2234), rabbit anti-EGFR (#4267), rabbit anti-phospho-ERK1/2 (Thr202/Tyr204, #4370), rabbit anti-ERK1/2 (#4695), rabbit anti-GAPDH (#5174) were from Cell Signaling. Rabbit anti-hB3GALT6 antibody (#55049-1-AP) was from ProteinTech, rabbit anti-hSLC39A9 antibody (#PA5-75557) was from Invitrogen and rabbit anti-EMP1 antibody (#bs-0558r) antibody was from Bioss Antibodies. Cross-adsorbed goat anti-mouse IgG-HRP (#G21040), goat anti-rabbit IgG-HRP(#G21234) and goat anti-mouse IgM-Alexa568 (#A21043) were purchased from Invitrogen.

Dextran sulfate sodium salt (6.5-10 kDa, #441490050) was purchased from Scientific Chemicals. Heparan sulfate (#S5992) and chondroitin sulfate (#S2416) were from Selleck Chemicals LLC. Dextran (60-90 kDa, MP Biomedicals, # 0218014010), Dextran Sulfate Sodium Salt (36-50 kDa, MP Biomedicals, # 0216011001), Heparin sodium salt (MP Biomedicals, # 0219411450), and alginic acid sodium salt (High Viscosity, # 02154723) were from MP Biomedicals. Dextran (40kDa, MilliporeSigma, #31389), Dextran (450-650kDa,

MilliporeSigma, #31392), Dextran sulfate sodium salt (>500kDa, MilliporeSigma, #D6001), alpha-cyclodextrin (MilliporeSigma, # C46421G), alpha-cyclodextrin sulfated sodium salt (MilliporeSigma, #4945425G), and hyaluronic acid sodium salt (MilliporeSigma, #53747) were purchased from Millipore Sigma.

### Lentivirus preparation and host cell transduction

Lentiviral transductions were performed as previously described ^51–53^. Briefly, HEK-293T cells were transfected with the lentiviral packaging plasmids psPAX2 (Addgene, #12260) and pCMV-VSVG (Addgene, #8454), along with the appropriate cargo plasmid using the XtremeGene9 transfection reagent (MilliporeSigma, #30353-5182) in accordance with the manufacturer’s instructions. After 48 hours, the HEK-293T culture supernatant was collected, filtered through a 0.45 µm PVDF filter (Millipore, #slhv033rs), and added to target TR146 cells cultured to 70-80% confluency in 6-well plates. The TR146 cells were infected by adding polybrene (MilliporeSigma, #H9268) to the medium at a final concentration of 4 µg/mL and centrifuging the tissue culture plates at 1000 × g for 30-60 minutes at 30°C. Two days after infection, puromycin was added to medium and maintained for 7 days to select for transduced cells.

### Construction of Cas9-expression TR146 cells

The TR146 cells were transduced with lentivirus that carried the spCas9-Blast construct (Addgene, #52962) at MOI = 0.3 as described above. Two days post transduction, the TR146 cells were incubated in selective medium containing 5 μg/ml of blasticidin (Gibco, # A1113903) for 7 days. Cas9-expression in single cell derived colonies was confirmed by western blotting with an anti-Cas9 antibody. A few clones were further tested in cell damage and uptake assay (Supplementary Fig. 1) to verify that the Cas9-expression TR146 cells were comparable to the wild type TR146 cells.

### Genome-wide CRISPR-Cas9 screening with candidalysin

The TR146 CRISPR genome-wide knockout library was generated using the Brunello human whole genome sgRNA library (Addgene, #73178) ^18,54,55^. This library contains a total of 76,441 sgRNAs with 4 sgRNAs per gene and 1000 control sgRNAs. The sgRNA library was transduced into TR146-Cas9 cells at MOI ∼0.3. For selection of transduced cells, puromycin (Thermo Scientific, # J672368EQ) was added to the medium at a final concentration of 0.4 μg/ml. A total of 60 million cells were transduced, ensuring that each sgRNA was represented ∼200 times. For screening, the transduced cells were exposed to 30 μM of synthetic candidalysin (Biomatik, de novo synthesized) in serum free DMEM/F12. Control cells incubated in medium alone were processed in parallel. After 8 h of candidalysin exposure, the surviving cells were rinsed and passaged twice in fresh candidalysin-free medium. This process was repeated five times on 2 separate occasions. The cells from last round were expanded and harvested, after which their genomic DNA was extracted using the Quick-DNA Midiprep Plus Kit (ZYMO Research, #D4075). DNA fragments containing the sgRNA sequences were amplified by PCR using primers lentiGuidePCR1-F (AATGGACTATCATATGCTTACCGTAACTTGAAAGTATTTCG) and lentiGuidePCR1-R (GTCTGTTGCTATTATGTCTACTATTCTTTCCCCTGCACTG) and purified with CleanNGS SPRI Beads (Bulldog Bio, #CNGS050) according to the manufacturer’s instructions. Next-generation sequencing (Illumina HiSeq) was performed by Novogene. The sequencing results were analyzed on the web platform PinAPL-Py (http://pinapl-py.ucsd.edu/) ^20^. The original sequencing data were deposited to the NCBI under BioProject PRJNA1081917.

### Construction of TR146 cells with targeted gene disruptions

All sgRNA oligonucleotides (Supplementary Table 2) were obtained from MilliporeSigma and cloned into the pLentiGuide-Puro plasmid (Addgene, #53963) as previously described ^51,52^. Briefly, pLentiGuide-Puro was digested with BmsBI (Fermentas) and purified using the GeneJET gel extraction kit (ThermoScientific, #K0692). Subsequently, each pair of oligonucleotides was annealed and phosphorylated with T4 PNK (New England Biolabs, # M0201) following the manufacturer’s instructions. The annealed oligonucleotides were ligated with the BsmBI-digested pLentiGuide-Puro plasmid which was then validated by PCR and Sanger DNA sequencing. Each pLentiGuide-Puro containing sgRNA was packed into lentivirus and transduced into TR146-Cas9 cells as described above. After 1 week of selection with puromycin, disruption of the targeted gene was verified by immunoblotting with antibodies directed against the corresponding gene product.

### siRNA of *GBF1* and *TYK2*

TR146 cells grown in antibiotics-free media at ∼70% confluence in 6-well plate were transformed with lipofectamine RNAiMAX reagent (Invitrogen, #13778) according to the manufacture’s protocol. Twenty nanomolar *GBF1* siRNA (Santa Cruz Biotechnology, #SC-105388), tyk2 siRNA (Santa Cruz Biotechnology, #SC-36764) or control scramble siRNA (Santa Cruz Biotechnology, #SC-37007) was applied to TR146 cell twice (time 0 h and time 24 h). Forty-eight hours post transfection, the cells in supplement-free media were seeded into 48-well plate overnight for the cell damage assay (time 64 h post transfection). The total protein of transfected cells was also collected at 64h for western blotting to confirm the successful knockdown of Gbf1 and Tyk2.

### Cell survival assay

The susceptibility of the host cells to damage by candidalysin and live *C. albicans* was determined using an XTT assay. The night before the experiment, host cells were seeded into a 48-well tissue culture plates in supplement-free medium. The next day, the cells were rinsed with warm PBS and then incubated with varying concentrations of candidalysin in serum-free DMEM/F12 medium (for TR146 cells) or DMEM media (for A431 cells) for 6 h. When inhibitors were used, they were added to the host cells either immediately before the candidalysin when the GAGs and GAG analogs were tested or 1 h before the candidalysin when the Tyk2 inhibitors deucravacitinib (3 μM, Selleck Chemicals #S8879) and TCJL37 (1 μM, Tocris Bioscience #6125) were tested. All inhibitors remained in the tissue culture medium for the duration of the experiment and host cells exposed to the inhibitors in the absence of candidalysin were processed in parallel. At the end of the incubation period, the medium above the cells was aspirated and replaced with phenol red-free DMEM/F12 medium containing XTT (ATCC, #30-1011K). After a 60-min incubation, 150 μL of the supernatant was transferred to a clean 96-well plate for measuring OD_475nm_ and OD_660nm_. The percentage of cells surviving was calculated relative to cells that were not incubated with candidalysin. The percent protection by the inhibitors was calculated using the formula: 100 × (sample-candidalysin) / (100-candidalysin).

The susceptibility of the host cells to damage by *C. albicans* was determined similarly except that the host cells were infected with *C. albicans* at a MOI of 5 for TR146 cells. After 6 h of infection for TR146 cells, the medium was replaced with serum-free medium containing 2 μg caspofungin per mL and the cells were incubated in this medium for 2 h to kill the *C. albicans* cells prior to performing the XTT assay. All experiments were repeated three times, with 2-3 replicates each.

In some experiments, an LDH-Cytox assay (Biolegend, #426401) was used to measure epithelial cell damage. In these experiments, the epithelial cells were grown in a 96-well plate and processed according to the manufacturer’s directions. The spontaneous LDH release was determined using uninfected cells that were processed in parallel (Untreated). To determine the maximum LDH release (MaxLDH), uninfected cells were lysed with 0.5% Triton X-100. Cell damage was calculated as: 100 × (Treated-Untreated) / (MaxLDH-Untreated).

### Western blot analysis

TR146 cells were seeded into each well of a 24-well plate (2×10^5^ cells/mL) the night before the experiment. The next day, they were incubated with either 10 µM candidalysin or *C. albicans* at MOI = 5 for various times. After aspirating the medium above the cells, the epithelial cells were lysed with 2× Laemmli SDS-PAGE sample buffer and the cell lysates were then incubated at 95℃ for 5-10 min. The lysates were resolved by SDS-PAGE, transferred to Immobilon-P PVDF membranes using the wet transfer method, and probed with specified antibodies at 4°C for overnight. After 3, 10 min washes with PBST, the membrane was incubated with goat anti-mouse or goat anti-rabbit IgG-HRP secondary antibody in 5% fat-free milk at room temperature for 45-60 min. Antigen signals were detected using Radiance Plus substrate (Azure Biosystems, #AC2103) and were visualized and imaged using a C400 imager (Azure Biosystem).

### Surface plasmon resonance assays

The GAGs used were porcine intestinal heparin (16 kDa) and porcine intestinal heparan sulfate (12 kDa, Celsus Laboratories); chondroitin sulfate A (20 kDa) from porcine rib cartilage (MilliporeSigma), dermatan sulfate (also known as chondroitin sulfate B, 30 kDa, from porcine intestine, MilliporeSigma), chondroitin sulfate C (, 20 kDa, from shark cartilage, MilliporeSigma). *N*-desulfated heparin (14 kDa) and 2-*O*-desulfated heparin (13 kDa) were prepared as described previously ^56^. 6-*O*-desulfated heparin (13 kDa) was provided by Prof. Lianchun Wang (University of South Florida). Sensor SA chips were from Cytiva and surfaced plasmon resonance measurements were performed on a BIAcore 3000 operating with BIAcore 3000 control and BIAevaluation software (version 4.0.1) (Cytiva).

Heparin coated biosensor chip was prepared as previously described ^57^. Briefly, heparin was biotinylated by mixing 2 mg heparin and 2 mg amine-PEG3-Biotin (Thermo Scientific), followed by the addition of 10 mg NaCNBH_3_. After incubation at 70°C for 24 hours, another 10 mg NaCNBH_3_ was added, and the mixture was incubated at 70°C for another 24 h. The final product was desalted using a spin column (3000 molecular weight cut-off) and then lyophilized. The biotinylated heparin was immobilized on a streptavidin chip following the manufacturer’s protocol, where a 20 μL solution of the biotinylated heparin (0.1 mg/mL) in HBS-EP buffer (0.01 M HEPES, 0.15 M NaCl, 3 mM EDTA, 0.05% surfactant P20, pH 7.4) was injected over flow cell 2 (FC2) of the SA chip at a flow rate of 10 μL/min. Successful immobilization of heparin was confirmed by the observation of a ∼200 resonance unit (RU) increase in the sensor chip. The control flow cell (FC1) was prepared by 1-minute injection with saturated biotin in HBS-EP buffer.

To measure the kinetics of the candidalysin-heparin interaction, candidalysin samples were diluted in HBS-EP buffer. Different dilutions of candidalysin were injected at a flow rate of 30 µL/min. At the end of the sample injection, HBS-EP buffer was passed over the sensor surface to facilitate dissociation. After a 3-min dissociation time, the sensor surface was regenerated by injecting 30 µL of 0.25% SDS. The response (RU) was monitored as a function of time (sensorgram) at 25 °C.

To test the inhibition of the candidalysin-heparin interaction by other GAGs and chemically modified heparins, candidalysin at 5 µM was pre-mixed with 10 µg/mL of GAG or 1 µM of chemical modified heparin and injected over the heparin chip at a flow-rate of 30 µL/min. After each run, a dissociation period and regeneration protocol were performed as described above.

### Flow cytometry

To measure calcium flux, TR146 cells and *B3GAT3*^-/-^ cells at 1×10^6^ cells/ml were loaded with the calcium probe Indo-1/AM (4 µM, Biotium, #500431) for 30 min at 37°C in cell loading medium (CLM, DMEM/F12+ 2% heat-inactivated fetal bovine serum + 25 mM HEPES) containing 1 mM EGTA. Cells were washed twice and kept at 37 °C in CLM at a concentration of 1 × 10^6^ cells/mL for another 30 minutes before flow cytometry. Calcium flux measurements were performed on a FACSymphony™ A5 cytometer (BD Biosciences) as suggested by the manufacture. After a baseline Indo-1 fluorescence was recorded for 90 sec, cells were treated with 1 µM of calcium ionophore A23187 (Sigma Aldrich, #C7522) or 10 µM of candidalysin while cell acquisition continued. Acquisition was performed for additional 6 min after treatment. The calcium flux was calculated as the ratio of fluorescence intensity at 395 nm (Ca^2+^-bound) and 496 nm (Ca^2+^-free), and followed over time. A kinetic analysis was performed using FlowJo software and the normalized (to the baseline) smoothed means of fluorescence ratios were plotted.

To quantify the levels cell surface heparan sulfate, 1×10^6^ of TR146, *XYLT2*^-/-^, *B3GALT6*^-/-^ and *B3GAT3*^-/-^ cells were fixed with 4% of paraformaldehyde for 10 min, and then blocked with 5% goat serum for 30 min before a 1-h incubation with a primary anti-heparan sulfate antibody (clone F58-10E4, #3702551) at 1:100 in FACS buffer (PBS + 5% FBS). After washing 3 times with FACS buffer, cells were incubated with goat anti-mouse IgM-Alexa633 (#A21046) at 1:100 in FACS buffer for 1 h. Cell surface heparan sulfate was quantified using a FACSymphony™ A5 cytometer and the results were analyzed with FlowJo software.

### C-laurdan emission assay

C-laurdan (Tocris, # 7273) was dissolved in chloroform to 1.1025 mM (e = 12,200 M^-1^cm^-1^) and further diluted with 200 proof ethanol to create a 100 μM stock suspended in a 91% ethanol, 9% chloroform solution. Samples were prepared in an aqueous buffer (150 mM NaCl, 10 mM HEPES) and incubated with either dextran sulfate (>500 kDa, MilliporeSigma # D6001) or heparin (Tocris # 2812) prepared in ultrapure water. Samples with and without candidalysin were loaded into a black 96-well plate (Corning) with 1 μM C-laurdan. The emission spectra from 400-600 nm were read on a Cytation 5 plate reader (BioTek) using an excitation wavelength of 350 nm. Spectral blanks without dye were subtracted. GP values were calculated using the following equation: GP= (I_blue_-I_red_)/(I_blue_+I_red_), where I_blue_ is the summation of fluorescence intensity values within the emission range of 420-460 nm and I_red_ is the summation of fluorescence intensity values within the emission range of 470-510 nm.

### Fluorescent dye-release assay

Fluorescent dye-release experiments were performed as described previously ^4^. In brief, lipid films of POPC (Avanti Polar Lipids) were rehydrated with 50 mM calcein solubilized in 50 mM EDTA and 50 mM NaPi (pH 8). Large unilamellar vesicles (LUVs) were formed by extrusion through a 100 nm filter, purified using a PD-10 desalting column (GE Life Sciences), and diluted to a working concentration of 144 μM. GAGs were suspended in MilliQ H_2_O at 50 mg/mL and 5 mg/mL and mixed with LUVs prior to measurement. Candidalysin suspended at 0.72 μM in CL Buffer (150 mM NaCl and 10 mM HEPES) was incubated with GAG + LUVs at a lipid to peptide molar ratio of 200:1 at the time of measurement in a 96-well black plate. Fluorescent readings were taken on a Cytation 5 plate reader using an excitation wavelength of 495 nm and an emission wavelength of 515 nm. Measurements were collected for two h.

### Atomic force microscope imaging

Candidalysin was hydrated in deionized water to 100 µM. The stock solution was divided into 10 µL aliquots and stored at −80 °C. DOPC stock in chloroform (Avanti Polar Lipids, Alabaster, AL) was dried under argon gas and vacuum overnight. Lipid films were resuspended in imaging buffer (10 mM HEPES, 150 mM NaCl, pH 7.3) and extruded through 100 nm filters (Whatman, United Kingdom) to form LUVs using a mini-extruder (Avanti Polar Lipids, Alabaster, AL). The solution was aliquoted and stored at −80 °C. Lipid concentration was determined using a standard phosphorus assay^58^. For control experiments, a candidalysin aliquot was thawed and diluted to 333 nM in imaging buffer. The samples were incubated at 25°C for 30 min, then 90 µL was deposited onto freshly cleaved mica discs (Ted Pella) and incubated for an additional 10 min. Loosely bound particles were washed away via buffer exchange (90 µL of buffer exchanged across the sample 5-6 times) Control experiments were also conducted to visualize candidalysin pores in DOPC in the absence of GAGs. These samples were prepared in a manner that replicated the previous control incubation experiments. A candidalysin aliquot was thawed and diluted to 555 nM in imaging buffer and incubated at 25°C. After 20 min, candidalysin was mixed with DOPC such that the final concentrations were 333 nM candidalysin and 200 μM DOPC. The solution was incubated 10 min to allow candidalysin to interact with liposomes, then 90 μL was deposited on mica. Samples were incubated for another 10 min to allow lipid vesicles to rupture and form bilayers. The lipid samples were rinsed via buffer exchange as before. Images were collected in imaging buffer with biolever mini tips (Olympus, *k* ∼ 0.1 N/m, *f_o_* ∼ 30 kHz in fluid) on a commercial instrument using tapping mode (Cypher, Asylum Research). Tip-sample forces were kept below 100 pN to reduce the likelihood of protein deformation.

To image candidalysin in the presence of GAGs, dextran, dextran sulfate, or heparan sulfate were diluted from stock concentrations in the imaging buffer. Each solution was mixed with candidalysin such that the final carbohydrate concentration was 5 μg/mL) and the candidalysin concentration was 333 nM. The samples were incubated 30 min at 25 °C prior to depositing 90 µL on freshly cleaved mica and incubating for an additional 10 min. The sample was washed via buffer exchange, then imaged. When imaging candidalysin and GAGs with lipid, candidalysin and GAG stocks were mixed and incubated for 20 min. This solution was then added to DOPC such that the final concentrations were 333 nM candidalysin, 5000 ng/mL carbohydrate, and 200 µM DOPC. This mixture was incubated an additional 10 min, then 90 µL was deposited on freshly cleaved mica and incubated another 10 min to allow for lipid bilayer formation. The sample was rinsed via buffer exchange and imaged. Image flattening and analysis was performed using a commercial software package (Asylum Research, Inc.). Loop and pore counts were done manually.

### Transmission electron microscope imaging

Transmission electron microscopy was performed using 3 µM candidalysin samples incubated for 30 min in imaging buffer (10 mM HEPES, 150 mM NaCl, pH 7.3) at 25 °C. Five microliter droplets were then incubated on carbon grids (Electron Microscopy Sciences) for 2 min at 25 °C and negatively stained with uranyl acetate. For candidalysin in the presence of GAGs, samples contained 3 μM candidalysin and 50 μg/mL of dextran, dextran sulfate, or heparan sulfate and were incubated for 30 min at 25 °C. TEM grids for these samples were prepared as described ^4^. Grids were imaged at 120 kV (JEOL, JEM-1400).

### Immunofluorescence microscopy

Epithelial cells grown on 12 mm round glass coverslips (Chemglass Life Sciences, #CLS1763012) were incubated with candidalysin for various times and then fixed with 4% paraformaldehyde (PFA) in PBS for 10 min at room temperature. After the cells were blocked with 0.1% BSA or 5% goat serum in PBS for 1 h, they were incubated in primary antibody diluted in PBS + 0.1% BSA for 1 h. The coverslips were washed and then stained for 1 h with fluorophore-conjugated secondary antibody and/or DAPI (MilliporeSigma, #MBD0015) in the dark. They were mounted inverted onto glass slides using ProLong Gold Antifade Reagent (Cell Signaling, #9071) and sealed with nail polish before imaging with Olympus IX83 fluorescence microscope. The number of CFSE-labelled candidalysin aggregates were counted manually. The colocalization of fluoresces signals was analyzed with the built-in modules in the CellSens Olympus microscope software.

### Epithelial invasion assay

*C. albicans* invasion of epithelial cells was determined using a differential fluorescence assay as described previously ^53,59,60^. Briefly, TR146 epithelial cells were grown on glass coverslips the day prior to infection and then infected with *C. albicans* blastospores at a MOI of 1 for 90 min. The cells were rinsed in a standardized manner and then fixed with 4% paraformaldehyde. The non-endocytosed fungal cells were stained for 1 h with a rabbit anti-*Candida*-antibody labeled with Alexa568. After rinsing with PBS, epithelial cells were permeabilized for 15 min in 0.1% Triton X-100 in PBS and the endocytosed and non-endocytosed fungal cells were stained with a rabbit anti-*Candida* antibody labeled with Alexa488. The coverslips were rinsed with PBS, mounted inverted on microscopic slides and visualized using epifluorescence microscopy. The percentage of invading *C. albicans* cells was determined by dividing the number of internalized cells by the total number of fungal cells per high power field, and normalized to the controls. A minimum of 100 fungal cells were counted on each coverslip.

### Cytokine assays

Host cells were grown in 48-well plates were incubated with either 10 μM candidalysin or *C. albicans* at a MOI of 5 as in the cell survival experiments. After 6 h and 24 h for TR146 and A431 cells, respectively, the culture supernatant was collected and stored at - 20°C. At a later time, the concentration of IL-1β (R&D, #DY20105; DB, #557953), GM-CSF (BD, #555126) and CXCL8 (BD, #555244) in the conditioned medium was measured by ELISA according to the manufacturer’s instructions.

### Murine model of VVC

The murine model of VVC was performed as described previously with slight modifications ^61–63^. Groups (n=7) of 6-8-week-old female C57BL/6 and CD-1 mice were purchased from Charles River Laboratories and housed in isolator cages mounted on ventilated racks. Mice were administered 0.1 mg β-estradiol 17-valerate (MilliporeSigma, #E1631-1G) dissolved in sesame oil subcutaneously 3 days prior to vaginal lavage or challenge with *C. albicans*. Stationary-phase cultures of *C. albicans* strain SC5314 were washed and adjusted to 5×10^8^ cells/ml in sterile endotoxin-free PBS. Mice were intravaginally inoculated with 10 μL of cell suspension, generating an inoculum size of 5×10^6^ blastospores. Colitis grade dextran sulfate sodium salt (DSS) (#160110, MP biomedicals) was dissolved with sterile cell culture grade water to 5 mg/mL, then mixed with carboxymethylcellulose sodium salt (CMC, Millipore, #217274-250GM) to make a 3% CMC gel. Vehicle (VEH) was composed of unsupplemented 3% CMC gel. Mice were intravaginally administered 20 µL of 5 mg/mL DSS or VEH gel formulation every 24 h from day −1 until day 2 post-inoculation (p.i.). At day 3 p.i., mice were sacrificed and vaginal lavage fluids (VLF) were obtained by flushing the vaginal canal with 100 μL of PBS.

### Assessment of fungal burden

Recovered VLFs were spiked with 1X final cOmpleteTM mini EDTA-free protease inhibitor (Roche, #11836170001). Fresh VLF (10 μL) was smeared onto Tissue Path Superfrost Plus Gold slides (Fisher Scientific), air dried, fixed with CytoPrep fixative (Fisher Scientific), and stored at room temperature. VLF were centrifuged at 4000 rpm, and supernatant transferred for storage at −80°C. Cell pellets were resuspended in their original volume of PBS as assessed by weight. An aliquot was serially diluted, plated on YPD agar plates containing 50 µg/ml chloramphenicol (#BP904-100, Fisher Scientific), and subsequently incubated at 30°C for 48 h ^64^. The resulting colonies were enumerated and log transformed as a measure of fungal burden. The Papanicolaou technique was used to assess polymorphonuclear leukocyte (PMN) recruitment by manually counting 5 nonadjacent fields by standard light microscopy using a 40X objective and reported as the mean. The cytokine IL-1β was measured by ELISA (Invitrogen, #88-7013A-88) according to manufacturer’s instruction. The ToxiLight™ Non-Destructive Cytotoxicity BioAssay (Lonza, #186467,) was used to measure adenylate kinase release in vaginal lavage fluid as a measure of tissue damage per the manufacturer’s instructions. Luminescence values were quantitated using a Synergy H1 microplate reader (Biotek).

## Supporting information

Supplemental table 1-CRISPR screen result

Supplemental table 2-sgRNA primers for knockcout

## Ethics statement

The animals used in this study were housed in AALAC-approved facilities located in the Regional Biocontainment Laboratory (RBL) at the University of Tennessee Health Science Center (UTHSC). The animal work conducted in this study was approved by the UTHSC Institutional Animal Care and Use Committee under protocol 21-0265 in accordance with the Guide for the Care and Use of Laboratory Animals. Every effort was taken to ensure that the use of animals was necessary for hypothesis testing, that the minimum number of animals required was utilized, and that steps were taken to minimize discomfort. Mice were given standard rodent chow and water ad libitum and monitored for signs of distress, including noticeable weight loss and lethargy.

## Data availability

The original sequencing data were deposited to the NCBI under BioProject PRJNA1081917. All data from this study are presented in the manuscript and supplementary figures and tables.

## Acknowledgements

This work was supported in part by grants R01DE026600 (to SGF), R01AI134796 (to BMP), R35GM140846 (to FNB), U01-AI124319 and U19-AI172713 (to MRY), S10OD028523 and R21AI156573 (to RJL and FZ) from the National Institutes of Health, USA; and 2122027 (to GMK) from the National Science Foundation, USA. JM received financial support from the China Scholarship Council award 201906150153 and the UTHSC Center for Pediatric Experimental Therapeutics. KGS acknowledges support from the Research Excellence Program at the University of Missouri. CMR was supported by a Graduate Advancement & Training Education Fellowship from the University of Tennessee-Oak Ridge Innovation Institute. We thank Zi-Qi Koo and Dr. Yu-Huan Tsai at Institute of Microbiology and Immunology in Taipei for the detailed protocol for labeling candidalysin with CFSE. The funding agencies had no role in the study design, data collection, data interpretation, or preparation of the manuscript.

---

**Supplementary Fig. 1.**
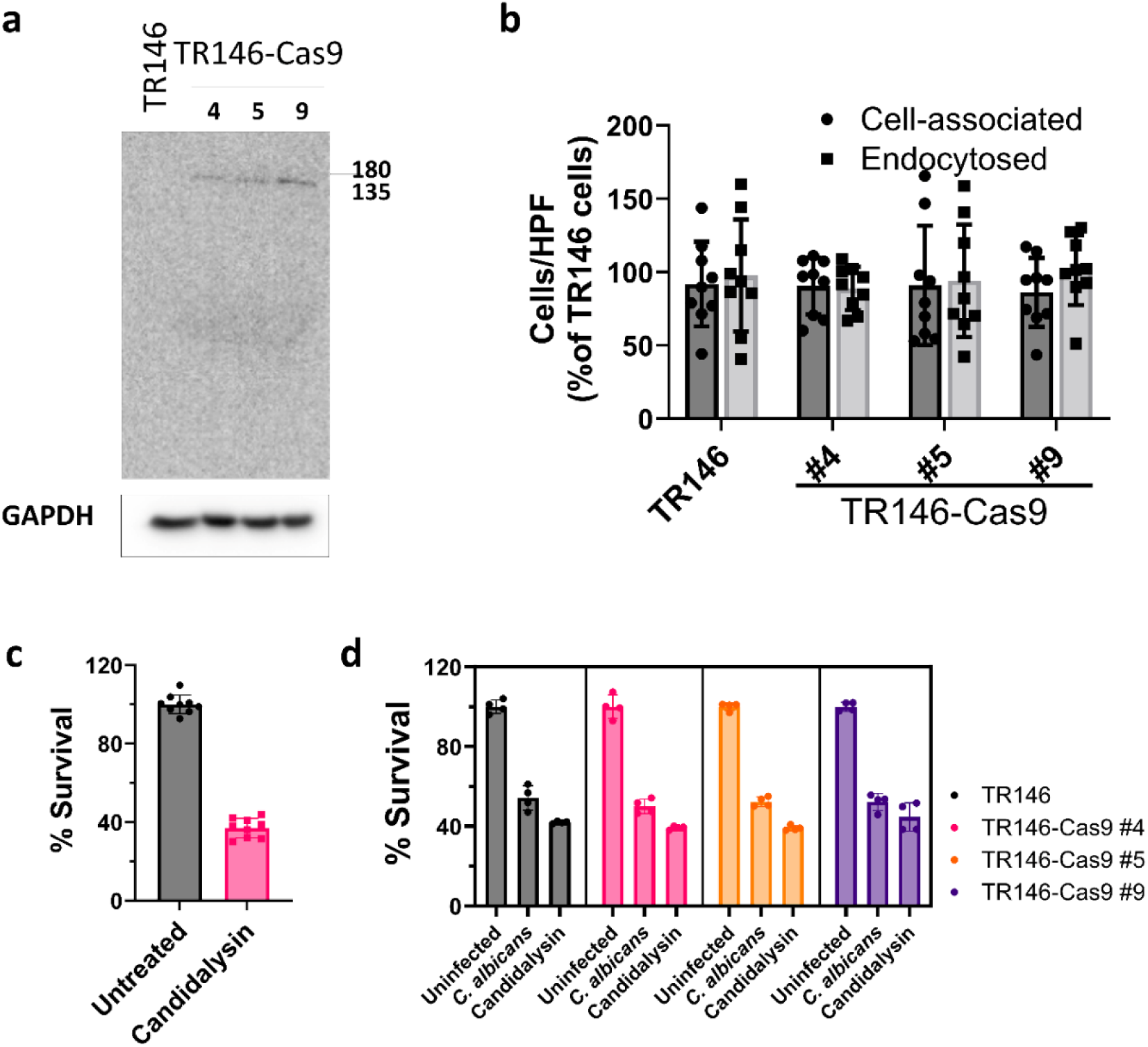
Cas9-expressing TR146 cells display comparable phenotypes to wild-type TR146 cells in response to wild-type *C. albicans* SC5314 and candidalysin. **a,** Western blotting of 3 clones of TR146 cells that stably expressed the P*_EF-1α_*-*Cas9*-*Blasticidin* construct. **b**, *C. albicans* association to and endocytosis by wild-type TR146 cells and the indicated clones of TR146-Cas9 cells. **c**, Cell survival of TR146 cells in response to candidalysin (30 μM) for 6 h as measured by an XTT assay. **d**, Survival of TR146 cells and TR146-Cas9 clones as measured by an XTT assay after a 6-h exposure to *C. albicans* (multiplicity of infection [MOI]=5) or candidalysin (30 μM). Clone 4 was selected for use in the subsequent experiments. Results are mean ± SD of 3 (**b, c**) or 2 experiments (**d**), each performed in triplicate (**b, c**) or duplicate (**d**).

**Supplementary Fig. 2.**
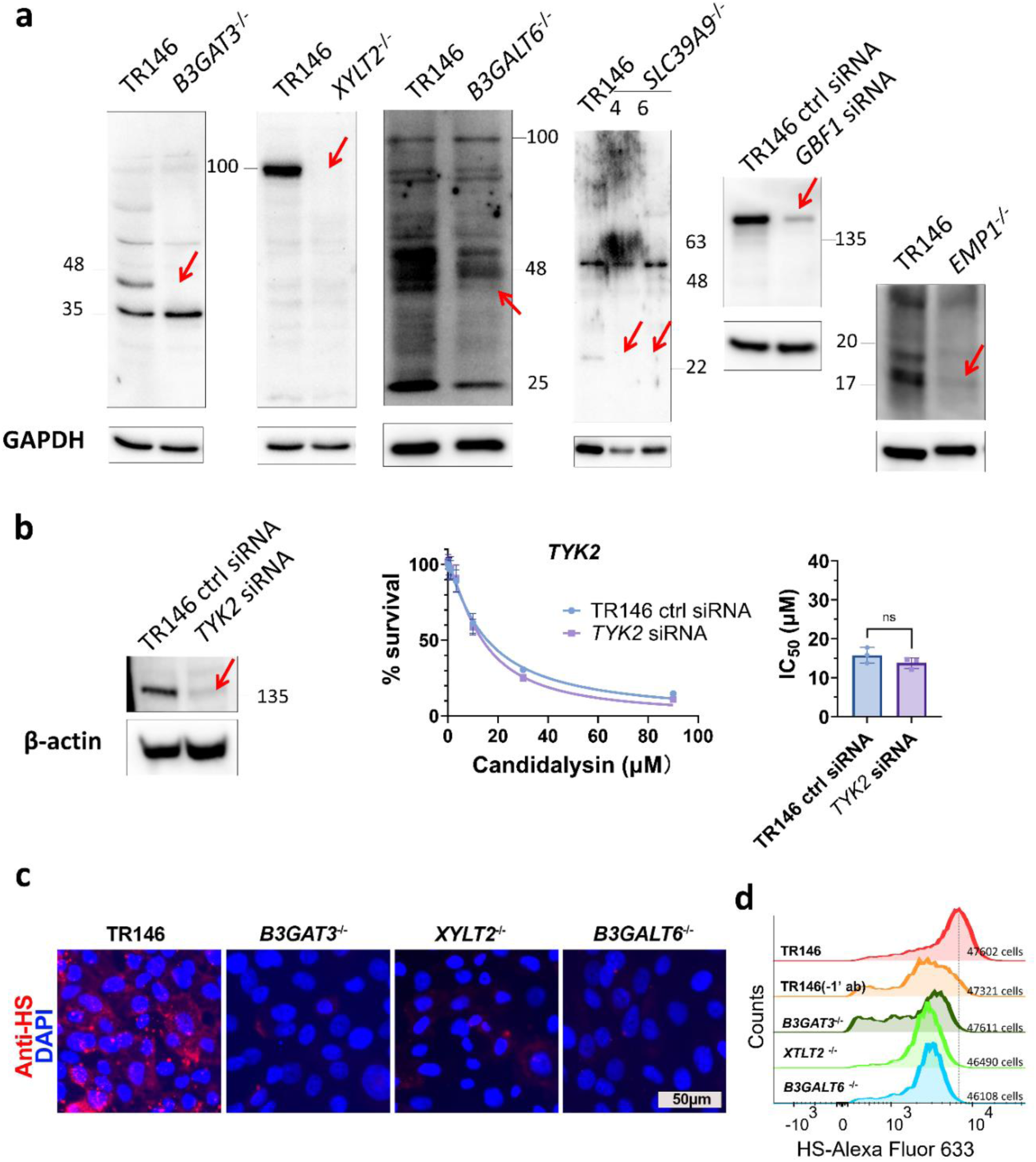
Western blots and immunofluorescence images of gene knockouts and knockdowns. **a,** Western blotting of B3gat3, Xylt2, B3galt6, Slc39a9 and Gbf1 in the corresponding CRISPR knockout cells or siRNA knockdown cells. Arrows indicate the target proteins. **b,** Tyk2 siRNA knockdown does not affect survival of oral epithelial cells after 6-h of candidalysin exposure. The left panel shows a representative Tyk2 western blot, and the middle panel shows the survival (measured by an XTT assay) of epithelial cells exposed to the indicated concentrations of candidalysin. The plots represent the combined results of 3 experiments, each performed in triplicate. The right panels show the concentration of candidalysin that yielded 50% survival (IC_50_), which was calculated from the data in the corresponding graph in the middle panel. Results are mean ± SD. ns, not significant by the unpaired, two-sided Student’s t test. **c**, Representative immunofluorescence images of TR146, *B3GAT3*^-/-^, *XYLT2*^-/-^ and *B3GALT6*^-/-^ cells stained with an anti-heparan sulfate antibody (red) and DAPI (blue). Scale bar: 50μm. **d**, Flow cytometric analysis of heparan sulfate expression on the surface of TR146 cells and the indicated mutants.

**Supplementary Fig. 3.**
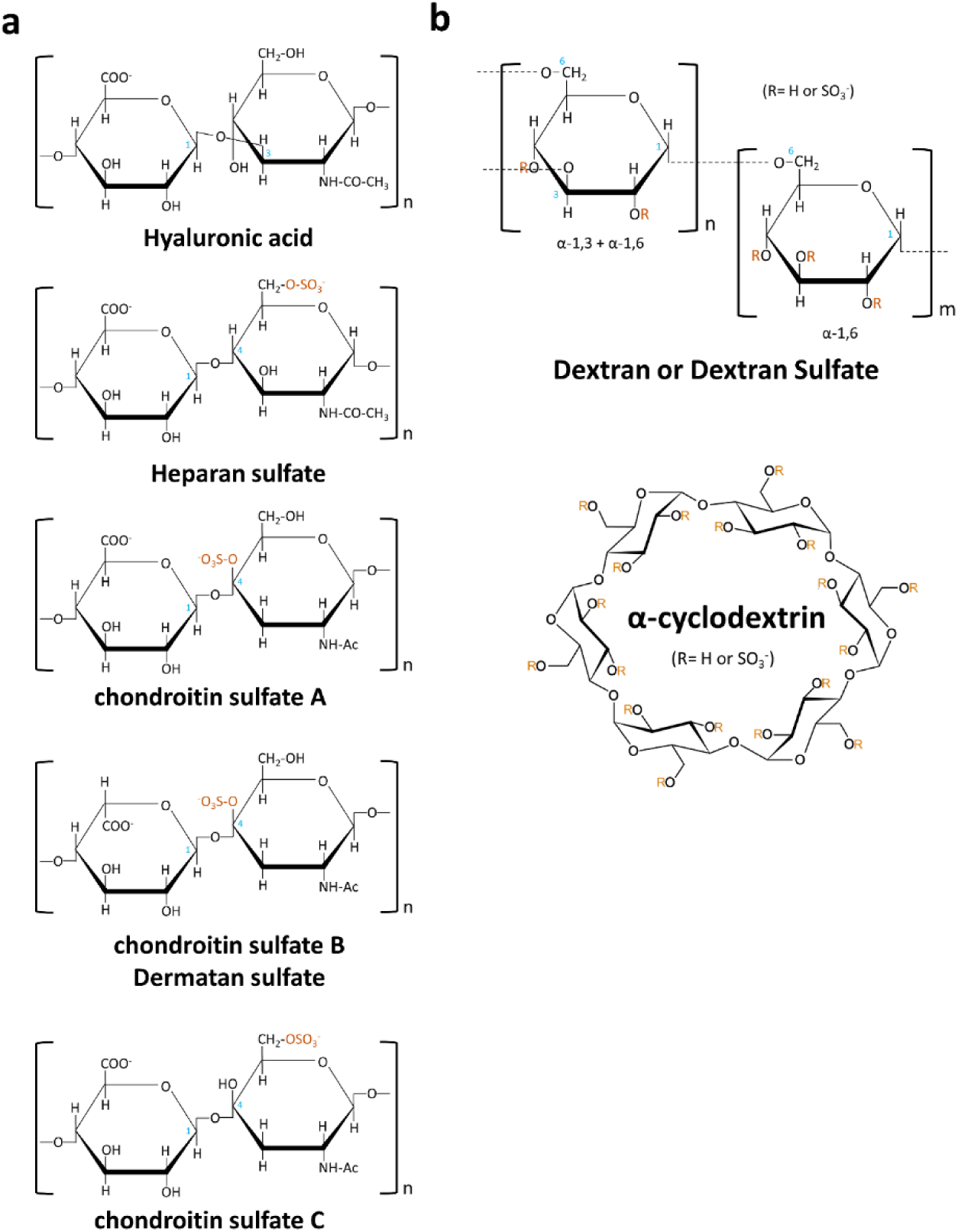
Structures of GAGs and GAG analogs used in the experiments. **a,** Structure of the naturally occurring GAGS, hyaluronic acid, heparan sulfate, chondroitin sulfate A, chondroitin sulfate B (dermatan sulfate) and chondroitin sulfate C. **b,** Structure of the GAG analogs dextran/dextran sulfate, and alpha-cyclodextrin/sulfated alpha-cyclodextrin.

**Supplementary Fig. 4.**
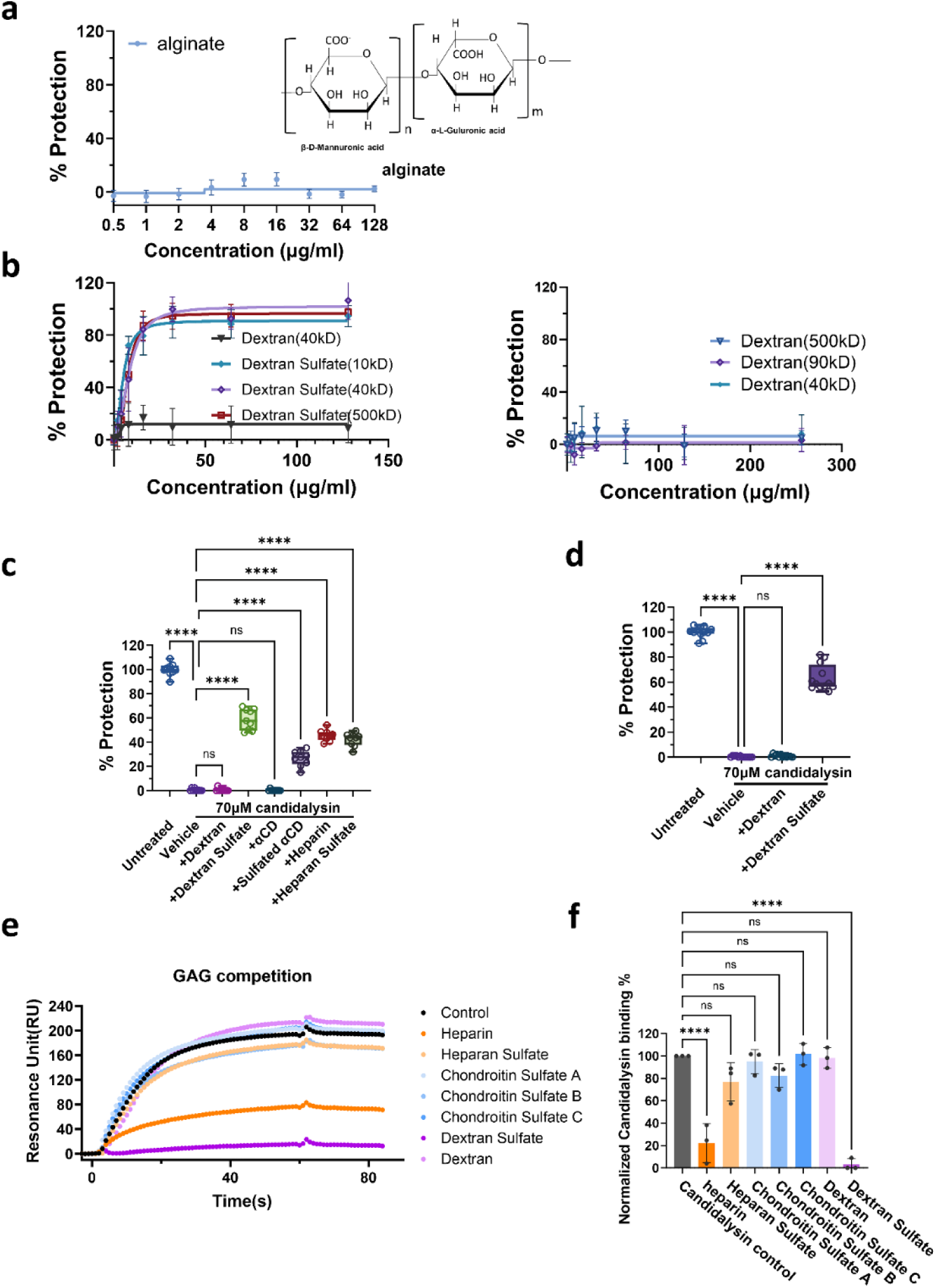
Sulfated GAGs but not carboxylate or non-sulfated GAGs protect epithelial cells from candidalysin-induced damage. **a,** Protection of oral epithelial cells from damage caused by a 6-h exposure to 30 μM candidalysin provided by alginate. **b,** Protection from damage caused by a 6-h exposure to 30 μM candidalysin provided by dextran and dextran sulfate (10kD, 40kD and 500kD) (left) or dextran (40kD, 90kD and 500kD) (right). **c**, Protection from damage caused by a 6-h exposure to 70 μM candidalysin provided by 100 μg/ml of dextran, dextran sulfate, α--cyclodextrin, sulfated α-cyclodextrin, heparin, and heparan sulfate. **d**, Protection from damage caused by a 24-h exposure to 70 μM candidalysin provided by 100 μg/ml of dextran, and dextran sulfate. **e,** Representative surface plasmon resonance sensorgrams showing the effects of heparin, heparan sulfate, chondroitin sulfate A, chondroitin sulfate B, chondroitin sulfate C, dextran and dextran sulfate on the interaction of candidalysin with heparin on a biosensor chip. **f,** Combined results of 3 independent experiments showing the inhibitory effects of the various GAGs or GAG analogs on the interaction of candidalysin with heparin. Results in **a-d** are mean ± SD of 3 independent experiments, each performed in triplicate. Protection was determined using an XTT assay. ns, not significant; ****, *p*<0.0001 by one-way ANOVA with Dunnett’s multiple comparisons test (**c**, **d**, and **f**).

**Supplementary Fig.5.**
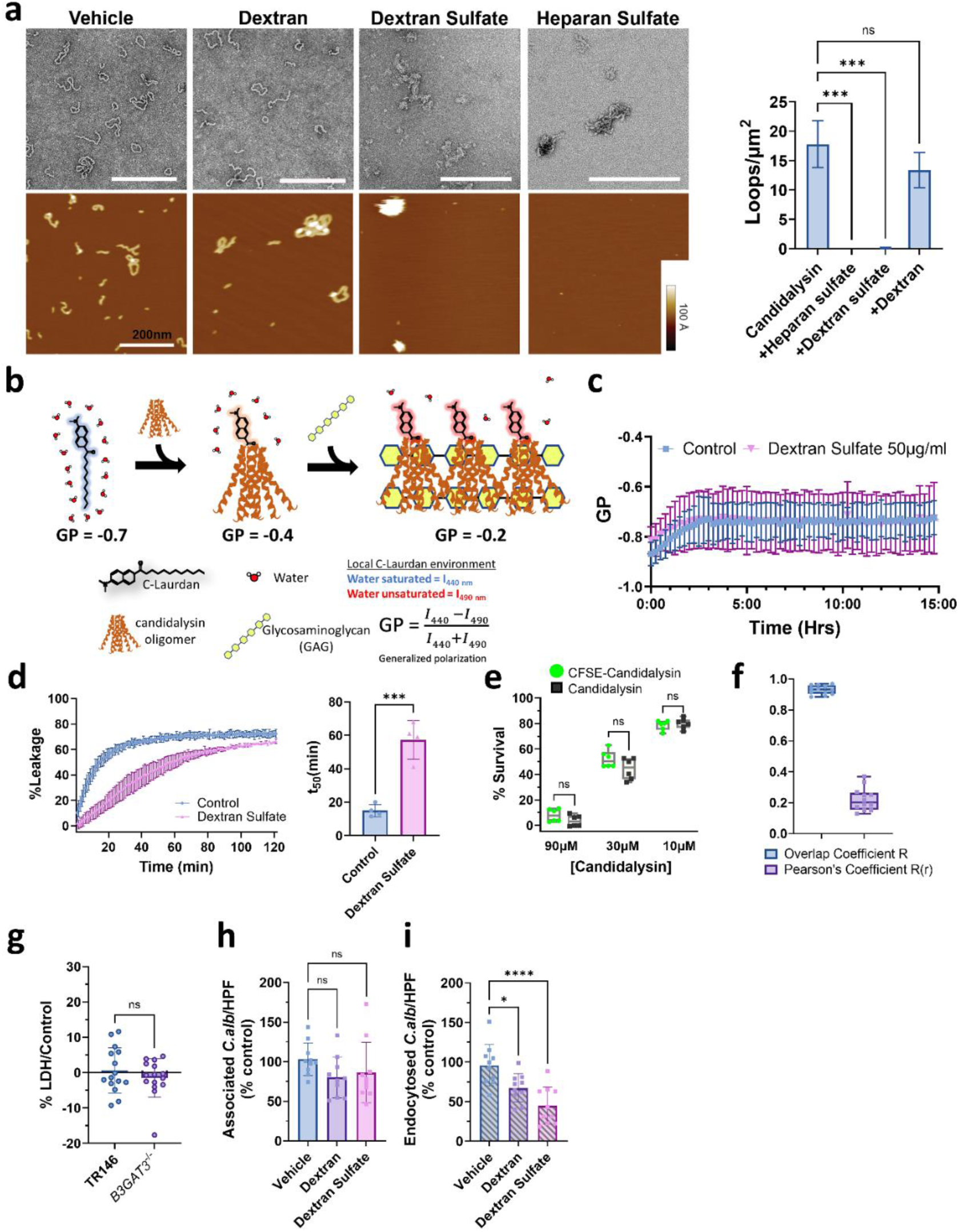
Candidalysin interacts with GAGs and their analogs. **a,** Transmission electron microscopy (TEM, top left) and atomic force microcopy (AFM, bottom left) images of candidalysin on a solid substrate with or without dextran, dextran sulfate and heparan sulfate. Candidalysin (3 µM) was incubated with dextran, dextran sulfate, or heparan sulfate (50 μg/mL) for 30 min before TEM imaging, and candidalysin (333 nM) was incubated with dextran, dextran sulfate, or heparan sulfate (5 μg/mL) for AFM imaging. The right panel shows the quantification of the AFM images in terms of loops per μm^2^ for each treatment. Scale bar: 200 nm. **b,** Diagram illustrating the C-laurdan assay. **c,** Time course of the effects of dextran sulfate (50μg/mL) on the GP score in the C-laurdan assay in the absence of candidalysin. Data are the mean ± SD of 3 experiments. **d,** Dextran sulfate (50 μg/mL) reduces the rate of candidalysin-induced membrane damage of 1-palmitoyl-2-oleoyl-glycero-3-phosphocholine (POPC) unilamellar vesicles. Representative time course (left). Combined results from 4 experiments (right). t_50_ values represent the time at which 50% membrane leakage occurred. Results are mean ± SD. The data were analyzed using an unpaired, two-sided Student’s test with significant *p*-values shown in comparison to the control. **e,** Comparison of the effect of candidalysin and CFSE-labeled candidalysin on the survival of oral epithelial cells after a 6-h exposure to 10, 30 and 90 μM candidalysin. Survival was measured using an XTT assay. Results are mean ± SD of two experiments, each performed in triplicate. **f,** Colocalization analysis of heparan sulfate and CFSE-candidalysin on TR146 cells. Overlap coefficient R and Pearson’s coefficient R(r) were generated by the Olympus microscopy software CellSens. Results are median (min to max) of 3 independent experiments, each quantifying 4 independent images. **g,** Damage to wild-type TR146 and *B3GAT3*^-/-^ cells caused by the *C. albicans ece1*Δ/Δ mutant (MOI = 5) after 5 h of infection, as measured by an LDH assay. Results are mean ± SD of three experiments, each performed in triplicate. **h, i**, Effects of 100 μg/ml dextran and dextran sulfate on the number of cell-associated and endocytosed cells of the *C. albicans ece1*Δ/Δ mutant. The average number of organisms per high-power field that were associated with and endocytosed by TR146 cells were 8.81 ± 1.81 and 2.27 ± 1.31, respectively. Results are mean ± SD of three independent experiments, each performed with a single replicate (**a, c, d**) or in triplicate (**e-i**). ns, not significant, \**p* < 0.05, \*\*\*\**p*<0.001 by unpaired, two-sided Student’s t-test (**d, g**) and one way ANOVA with Dunnett’s test for multiple comparisons (**h, i**).

**Supplementary Fig.6.**
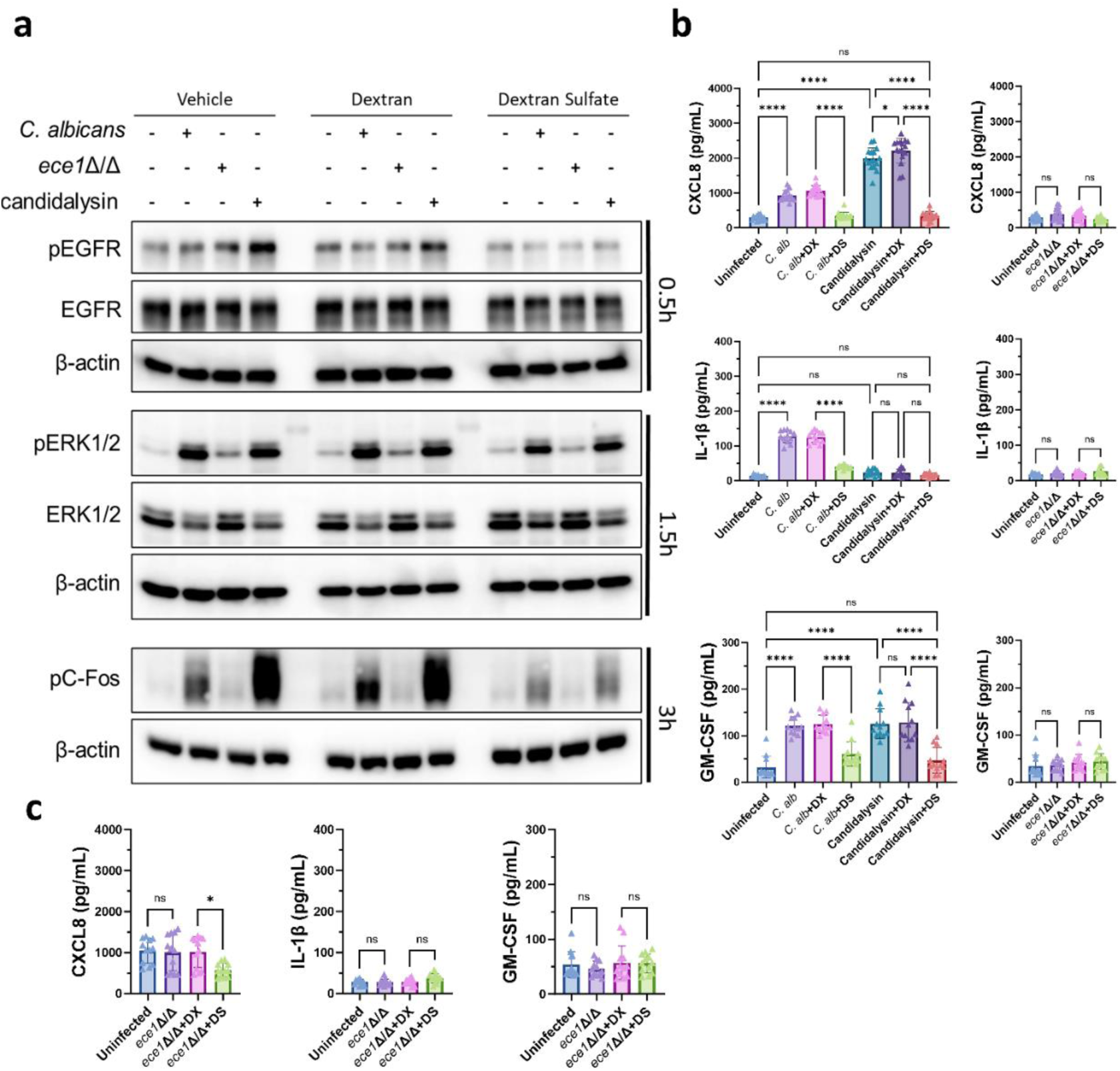
Effects of dextran and dextran sulfate on stimulation of *B3GAT3*^-/-^ cells by candidalysin and live *C. albicans*. **a**, Western blot results of epidermal growth factor receptor (EGFR), extracellular regulated kinase1/2 (ERK1/2) and the c-Fos transcription factor in *B3GAT3*^-/-^ epithelial cells induced by wild-type *C. albicans* SC5314 or the *ece1*Δ/Δ mutant (MOI = 5), or candidalysin (10 μM) with or without dextran (DX) and dextran sulfate (DS) (100 µg/ml) at the indicated time points. Shown are representative results of 3 independent experiments. **b**, The effects of dextran (DX) and dextran sulfate (DS) (100 μg/ml) on the production of CXCL8 (top), IL-1β (middle) and GM-CSF (bottom) by *B3GAT3*^-/-^ cells incubated for 6 h with wild-type *C. albicans* (MOI=5) or candidalysin (10 μM) (left panel) or infected with the *C. albicans ece1*Δ/Δ mutant (right panel). Results are mean ± SD of 3 experiments, each performed in duplicate or triplicate. **c**, Effects of dextran and dextran sulfate (100 μg/ml) on the production of CXCL8, IL-1β, and GM-CSF by TR146 cells infected with the *ece1*Δ/Δ mutant (right, MOI=5) for 6 hours. **, p < 0.01; ****, p < 0.0001 by one-way ANOVA with Dunnett’s multiple comparisons test (**b-c**).

**Supplementary Fig. 7.**
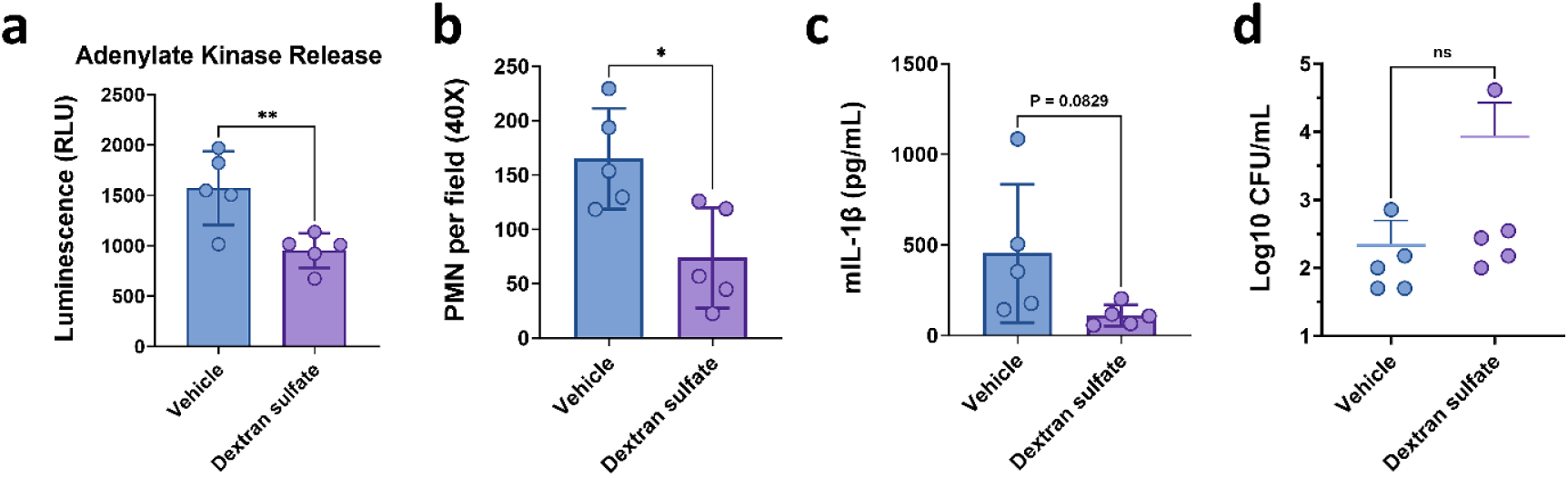
Dextran sulfate protects vaginal epithelial cells from damage and inhibits pro-inflammatory cytokine production in CD-1 mice with vulvovaginal candidiasis. **a-d,** CD-1 mice were treated with either dextran sulfate or vehicle alone intravaginally prior to vaginal inoculation with *C. albicans* SC5314 and daily thereafter. After 3 days of infection, the concentration of adenylate kinase (a measure of host cell damage) (**a**), neutrophils (PMN) (**b**), IL-1β (**c**), and fungal colony forming units (CFU) (**d**) in the vaginal lavage fluid was determined. Results in (**a-d**) are the mean ± SD of 5 mice per experimental group in a single experiment. ns, not significant; *, *p* < 0.05; **, *p* < 0.01; ***, *p* < 0.001; ****, *p* < 0.0001 by the student’s t-test (**a-d**).

**Supplemental Table 1.** Genes identified in the CRISPR screen.

**Supplemental Table 2.** Sequences of oligonucleotides used for guide RNAs

**Supplemental Table 3.** Experimental data.

